# Disordered information processing dynamics in experimental epilepsy

**DOI:** 10.1101/2021.02.11.430768

**Authors:** Wesley Clawson, Tanguy Madec, Antoine Ghestem, Pascale P Quilichini, Demian Battaglia, Christophe Bernard

## Abstract

Neurological disorders share common high-level alterations, such as cognitive deficits, anxiety, and depression. This raises the possibility of fundamental alterations in the way information conveyed by neural firing is maintained and dispatched in the diseased brain. Using experimental epilepsy as a model of neurological disorder we tested the hypothesis of altered information processing, analyzing how neurons in the hippocampus and the entorhinal cortex store and exchange information during slow and theta oscillations. We equate the storage and sharing of information to low level, or primitive, information processing at the algorithmic level, the theoretical intermediate level between structure and function. We find that these low-level processes are organized into substates during brain states marked by theta and slow oscillations. Their internal composition and organization through time are disrupted in epilepsy, loosing brain state-specificity, and shifting towards a regime of disorder in a brain region dependent manner. We propose that the alteration of information processing at an algorithmic level may be a mechanism behind the emergent and widespread co-morbidities associated with epilepsy, and perhaps other disorders.

## Introduction

Most, if not all, neurological pathologies, including Alzheimer’s disease, epilepsies, and Parkinson’s disease, aside from their specificities, display commonalities in terms of cognitive (e.g., memory) and mental (e.g., anxiety and depression) disorders (Hesdorffer, 2016). Historically, attempts have been made to correlate higher-level changes to the underlying structural alterations. However, structural alterations may be very different from one pathology to the next, even within a given brain disorder. The origin of shared and generic deficits must therefore be sought for at a level higher than the structural one. We hypothesize that diverse pathological mechanisms can lead to similar modifications of information processing, emerging from, and existing between, structural and functional levels. Whether information processing is modified in a pathological context is not known. Furthermore, a formal framework for the quantification of these processes is missing.

As a model of neurological disorder, we consider Temporal Lobe Epilepsy (TLE), the most common form of epilepsy in adults (Tatum, 2012). TLE is itself highly heterogenous in terms of differences of histopathology (Blumcke et al., 2013), semiology (Barba et al., 2007; Bartolomei et al., 2008) and cognition and mental state (de Barros Lourenco et al., 2020; Holmes, 2015; Krishnan, 2020). Such heterogeneity is also found in experimental models of TLE (Rusina et al., 2021). Structural alterations may change several features that are relevant for information processing, such as rate coding, temporal coding, synaptic plasticity, and network oscillations (Lenck-Santini & Scott, 2015). In keeping with this proposal, hippocampal place cells are unstable, firing becomes randomized during ripples, synaptic plasticity, and oscillations are altered, and these changes are correlated with deficits in hippocampus-dependent spatial memory in experimental epilepsy (Chauvière et al., 2009; Inostroza et al., 2013; Lenck-Santini & Holmes, 2008; Lopez-Pigozzi et al., 2016; Suarez et al., 2012; Valero et al., 2017). Given this diversity of deficits, it is reasonable to presume that in TLE local information processing is altered at a more fundamental level, with widespread impacts on multiple functions.

It is difficult to link specific alterations at the structural level to high order cognitive deficits as we do not know where information processing is localized, what is being processed, nor how it is integrated into function. In other words, with reference to the notion of the *algorithmic* level introduced by Marr and Poggio (1977), we do not know what are the “algorithms” that bridge structure and function. The common axiomatic view is that neural information processing stems from the spatiotemporal organization of the firing of neurons. Information theory was designed to be agnostic to the content of information and thus provides useful metrics to track primitive, or fundamental, information processing operations (Shannon, 1948). Neuronal firing intrinsically carries information due to its statistical properties. Auto-correlations in firing actively maintain this information through time - active information *storage* (Lizier et al., 2012; Wibral et al., 2014), and cross-correlated firing between different neurons allows the sharing of this information between themselves (Kirst et al., 2016). Focusing on such basic operations allows investigation of how patterns of coordinated neural firing may translate into primitive low-level information processing (Clawson et al., 2019), akin to the algorithmic level. Here, we hypothesize that the key differences between control and epileptic networks are not only present at the structural level, but also at a more general and core algorithmic level of quantifiable primitive operations.

To test this hypothesis, a multilevel experimental approach is required (Scott et al., 2018). Multi-channel electrode recordings of neural populations provide such a dataset which spans two levels of analysis: the action potential at the neuronal level and oscillations at the population level. As neural computation is brain state dependent (Quilichini & Bernard, 2012), we consider the global brain states of theta (THE) and slow oscillations (SO), which can be recorded during anesthesia. Previous work in control animals demonstrate that neuronal activity patterns in the hippocampus and entorhinal cortex switch between different information processing substates (IPSs) (Clawson et al., 2019). An IPS corresponds to an epoch in which primitive operations of information storage and sharing in a local microcircuit remain temporally consistent. IPSs continuously switch from one IPS to another, similarly to what has been described at higher level of organization, such as the dynamics of resting state networks and EEG microstates (Calhoun et al., 2014; Van de Ville et al., 2010). In the control hippocampus and entorhinal cortex, the sequences of IPSs are complex, i.e. standing between order and disorder (Clawson et al., 2019).

Using an unbiased quantification of IPSs, we compare their properties and organization between control and experimental epilepsy conditions. We focus on the hippocampus and the entorhinal cortex, two major structures commonly affected in TLE (Curia et al., 2008). We find that IPS’ internal organization and switching dynamics, although not suppressed, shift toward a less structured and more random spatiotemporal organization in experimental epilepsy than in control. Such disruption of information processing at the algorithmic level itself could underly the general performance impairments in TLE.

## Results

### Design

We analyze the local field potentials (LFPs) and action potentials from individual neurons measured in the hippocampus (CA1) and medial entorhinal cortex (mEC) from control (n = 5) and experimental epilepsy (n = 6) rats under anesthesia (Figure 1A-B, see Methods for details). Unsupervised clustering of the spectral content of LFPs reveals that field activity continuously switches between two states: slow oscillations (SO, 0.5-3 Hz) and theta oscillations (THE, 3-6 Hz) (Figure 1B, S1). As previously reported in freely moving animals (Chauvière et al., 2009), THE power and peak frequency are decreased in CA1 in experimental epilepsy (Figure S1). Although, the peak frequencies of THE and SO are not modified in the mEC in epilepsy, their power is decreased (Figure S1). However, both frequency and power ratios between SO and THE are similar in control and epilepsy.

**Figure 1.**
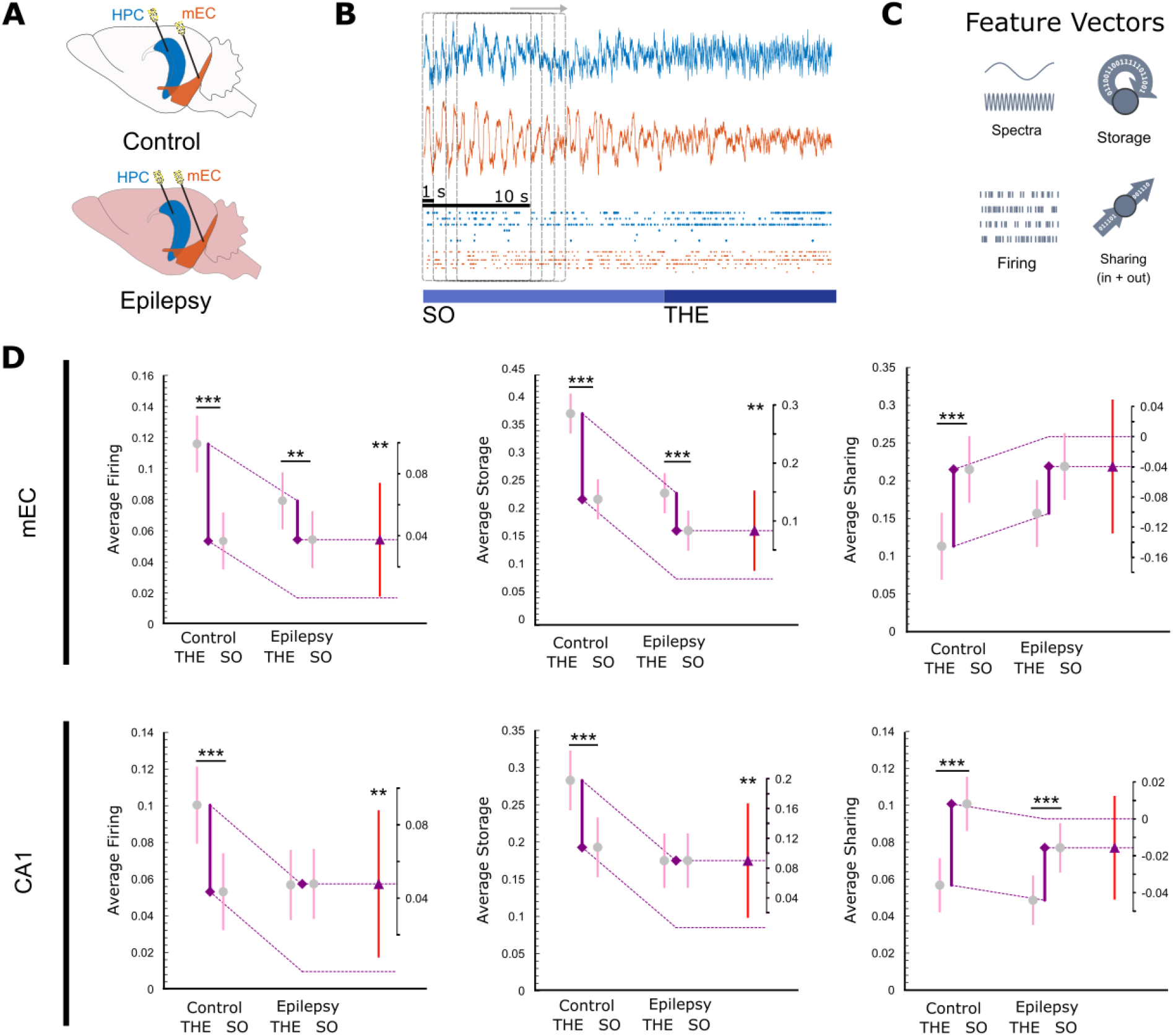
Experimental and analytical design. (A) Cartoon representing the approximate recording locations in mEC (orange) and CA1 (blue) in control and experimental epilepsy. (B) Example of LFP (top) and firing (bottom, each line represents one neuron, a dot represents an action potential) data recorded in control CA1 and mEC during SO and THE. Overlayed is a representation of our analytic method that uses 10 s long sliding windows shifted by 1 s at each step. (C) Cartoon examples of the four acquired data features. (D) Average values and difference of differences graphs for data features taken from spiking data during epochs of THE and SO in mEC (top) and CA1 (bottom) in both control and epilepsy conditions. See S2 for the same graph represented as a function of region, rather than oscillatory state. Circles and triangles represent the mean, and all bars represent a 99% bootstrapped confidence interval. Significance is shown using the symbol (*) with their standard corresponding meaning (*, p<0.05; **, p<0.01; ***, p<0.001). The numerical values are provided in Table S1.

We extract three features from the spike trains using a sliding widow procedure (Figure 1B-C): (1) firing, the number of times a neuron fired within a window, (2) storage, the information theoretical measure of active information storage (Lizier et al., 2012; Wibral et al., 2014), which captures temporal patterns of spiking for a single neuron within a window – notably in our case, how regular or repetitive these patterns are – and (3) sharing, an information theoretical measure of information sharing (Kirst et al., 2016), which captures spatiotemporal patterns of coordinated spiking across neurons within a window. First, we examine whether these features are dependent upon the brain state (THE versus SO), the region (CA1 versus mEC) and the condition (control versus epilepsy).

### Epilepsy reduces firing, storage and sharing differences between THE and SO states

In control animals, we find that in both regions, average firing and storage of all neurons is larger during THE than SO, while average sharing is lower (Figure 1D, see also S2), in keeping with the idea that neuronal computation is brain state-dependent (Quilichini & Bernard, 2012). In epilepsy, we find that average firing and storage are decreased during THE, but not during SO, as compared to control in both mEC and HPC. As a consequence, the brain state-dependency of firing and storage, which is consistent across controls, is reduced in both regions in epilepsy (Figure 1D). There is thus, in epilepsy, a large deviation from the operating mode found in control conditions.

We have previously shown that THE and SO states are in fact characterized by a complex dynamic organization in terms of firing, storage or sharing features (Clawson et al., 2019). A feature value (e.g., storage) can remain stable during a given time period (i.e., during successive windows), before switching to a different feature value with its own period of stability. We called these periods of stability *substates* of firing, storage or sharing. We begin by assessing the properties of substates in control and in epilepsy, as substate switching constitutes an important qualitative aspect of coordinated firing dynamics.

### Terminology, metrics, and methodology

Figure 2A illustrates an example of the procedure for a ~ 25 min long recording performed in the mEC in a control animal. Spectral analysis of the LFP reveals the alternation between THE and SO states (upper row). Through an unsupervised substate extraction procedure based on *k-*means clustering (see *Methods*), we identify in this example 4, 3, and 5 substates of stable patterns for firing, storage and sharing, respectively. The four features together, seen as 4 rows in Fig 2A, define a *switching table*. Each time point in the table corresponds to an *information processing state* (IPS), i.e. a combination of global state, firing rate, storage, and sharing patterns at this time point. By characterizing which neurons fire, how much, and with which correlation properties, an IPS provides a robust characterization of the pattern of coordinated activity occurring within each temporal window. Note that the switching transitions from one substate to the next are not necessarily synchronous between the different features, a property found in all recordings. In Figure 2B, we show, encoded as vertical color bars, the absolute values of firing, storage and sharing features that different neurons assume in the different substates. For a given feature, the values appear clearly different for a given neuron between substates. We will quantify these differences in the next section.

**Figure 2.**
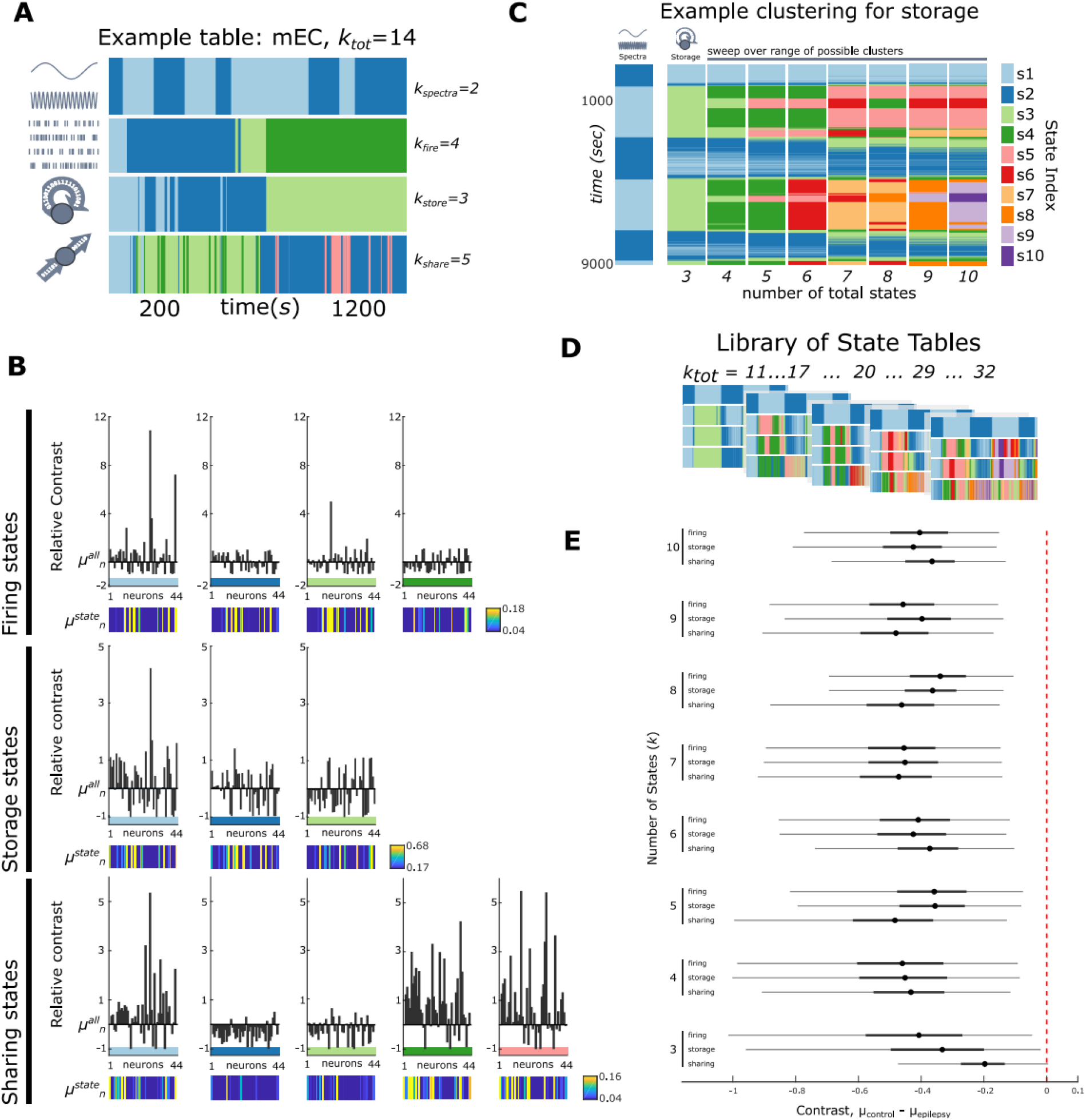
Clustering & contrast in control and epilepsy. (A) An example state table for the mEC in a control animal with a total state count of *k_tot_* = 14. The different substates are color coded. Note that switching is not synchronized across the different features. (B) Relative contrast values for the table given shown in (A). The substates shown in A are shown in B as a horizontal bar with the same color. Each graph shows the relative contrast of each of the 44 neurons, for each substate, and each feature. Below each graph is a visual indicator of a neuron’s feature values within the substate (vertical color bar). Here, the color scale varies from near 0, dark blue, to the top 10% of all average activity within the state. Therefore, any neuron whose activity is within this top 10% will be bright yellow. (C) Temporal dynamics (vertical axis) of storage substates as a function of *k* (horizontal axis). The far-left column shows the dynamics of THE and SO spectral states. (D) An example of a resulting state table library, or a collection of all possible combinations of all clustering with a range of *k_tot_ = 11 – 32.* (E) Average contrast difference between control and epilepsy is shown with respect to both feature and number of states, *k.* The circles represent the mean difference, the thick black bars represent the 25-75% quantile and the thin black bars represent the 1-99% quantile. The red dotted line is to add the null hypothesis line of no significant difference between control and epilepsy.

The switching table of Figure 2A is constructed using an unsupervised clustering algorithm, k-means, guided by an a priori assumption that (1) there exist separable clusters of data and (2) there are exactly *k* of these clusters (here 4, 3, and 5 for firing, storage and sharing, respectively). Using a null model, we demonstrate that there exist separable clusters (Figure S3). However, as the ground truth of how many clusters exist is unknown, statistical criteria can be used to find the optimal number (as done in Clawson et al., 2019). Here, we use a more general approach varying the *k* value for each firing, storage, and sharing feature while fixing *k* = 2 for the spectral feature. Each quadruplet of *k* values will produce a specific switching table. Figure 2C illustrates this concept, showing the resultant clustering of storage substates through time as *k* increases from 3 to 10. A low value may underestimate the real number of substates, while a large number may be an overestimate producing substates that rarely occur more than once (see Methods). We therefore use a lower bound of *k* = 3, and a reasonable upper bound of *k* = 10, wherein the clusters become too fine (Figure 2C, see Methods). We thus consider eight possible *k* values for each feature, giving rise to 8^3^ = 512 possible switching tables. Each switching table is characterized by the total number of substates it contains: *k_tot_* = 2 + *k_firing_* + *k_storage_* + *k_sharing_* with a maximum value of *k_max_* = 32 (32 = 2, the number of spectral states + 3 features × 10). The collection of all switching tables for a given recording defines a *library* of tables (Figure 2D). We chose such a method with the intention that without an a priori approach on the underlying principle, if we extract generic rules, they should be valid independently of the choice of number of clusters, at least for a reasonable wide range of *k* values. Now, all analysis that can be done on a switching table is performed for each library, which gives the added benefit of assessing the robustness of the results regarding the number of clusters.

### Substates are more contrasted in epilepsy

The vertical color bars in Figure 2B qualitatively show that individual neurons can take different firing, storage or sharing values across substates. In order to quantify these differences, we measure how “contrasted” are different substates. If we consider the firing feature of a given neuron, we first calculate its global mean firing rate (over the whole duration of the recording), and its mean firing rate within each substate. The relative contrast is defined as the difference between the substate mean firing rate and the global mean firing rate, normalized by the global mean firing rate. Evaluating contrast allows better tracking of the differing compositions of substates at the single neuron level. Figure 2B shows the relative contrast plots for the 44 recorded neurons and the various substates in the same dataset and substate decomposition we use as an example in Figure 2A. The differences between substates for each feature now clearly appear as large changes in the distributions of contrast values for the recorded neurons. Now, we extract the *substate contrast* of each substate for each feature - the average of the absolute values of the heights of the bars in the relative contrast plot. This substate contrast tells us how much a given substate stands out from its feature’s global average. Increasing the number of *k* substates may decrease the substate contrast.

Figure 2E shows the distributions of the differences in contrasts between control (n=5) and epilepsy (n=6), for firing, storage, and sharing features in the mEC, for the chosen *k* values (3 ≤ *k_firing_*, *k_storage_*, *k_sharing_* ≤ 10). For all values of *k*, for all features, the contrast differences lie entirely below zero, demonstrating that substate contrast is generally higher in epilepsy than in control. We also see no clear dependence upon *k* values, i.e., the number of substates. The same result is found in CA1, however higher bounds closer to the 99^th^ percentile do cross 0 (Fig S4). We thus identify another major alteration in epilepsy; substates are more contrasted, exhibiting more marked differences with respect to the mean. This suggest that in epilepsy, substate switching more strongly modulates the neural population with regards to firing, storage and sharing. While this seems to stand in contrast with the previously described reduction of the modulatory influence exerted by global oscillatory states, this may be explained by a disrupted articulation between substate and global state, as we explain in the following section.

### Loss of global state specificity of firing, sharing and storage substates in epilepsy

Since firing patterns are brain state-dependent, we assess whether this type of specificity is also found at the level of information processing substates. For a given state table in a library, we calculate the probability that a substate occurs during THE, SO or both. We name it *state specificity index* (SSI), a metric bounded between 0 (a substate occurs equally in THE or SO) and 1 (a substate is exclusive to either THE or SO) (see Methods). In control animals (Figure 3, blue curves), most substates are brain state specific in both mEC and CA1, independently of *k*. Most SSI values are above 0.8, well above the null hypothesis 0.23 ± 0.03 value of lack of global state specificity. Global state specificity of substates is thus a robust result in control animals with respect to *k*.

**Figure 3.**
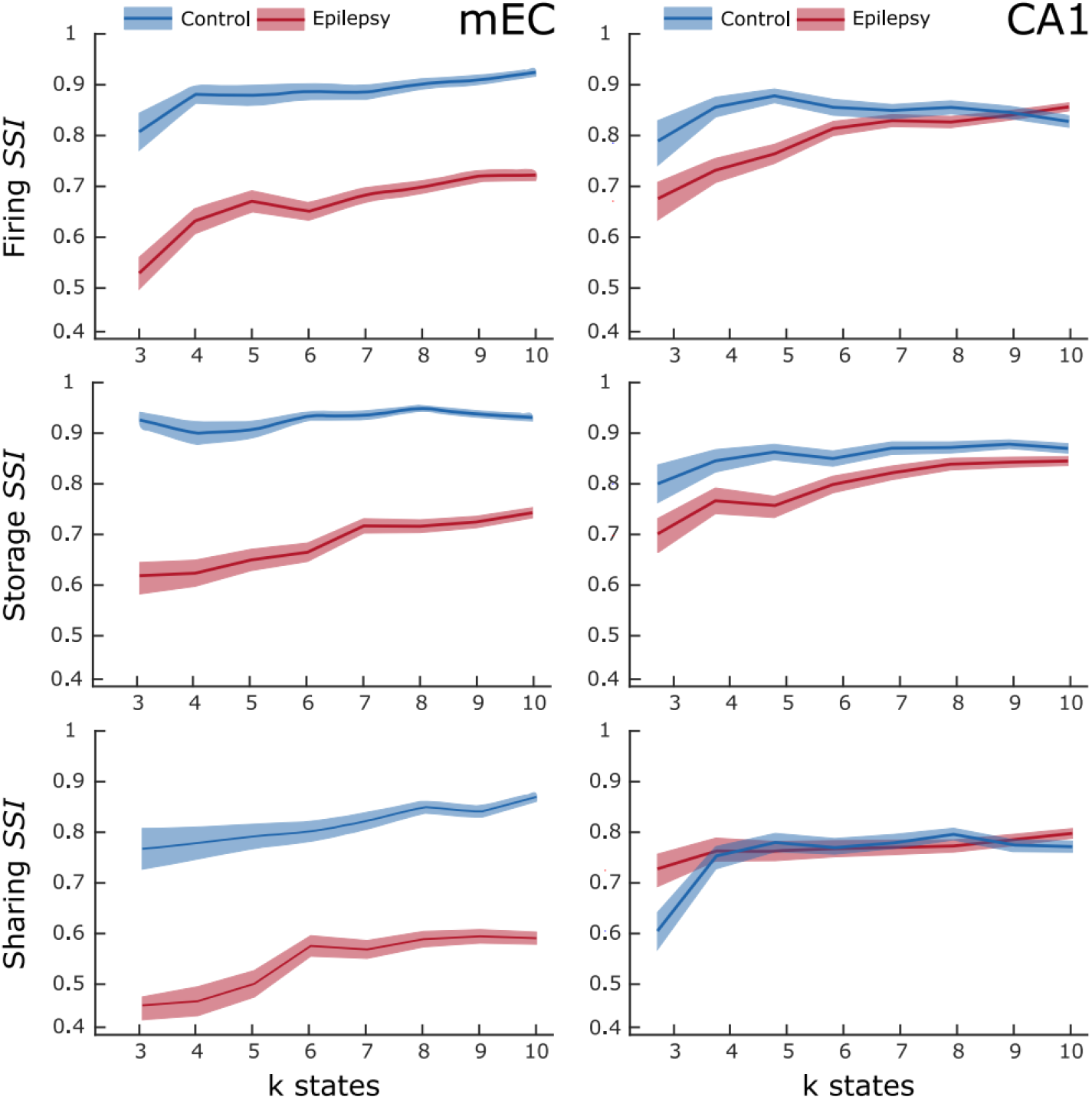
Loss of brain state-dependency of substates in the mEC in epilepsy. State similarity index (SSI) is shown here vs number of *k* states for each feature in mEC and CA1. Blue represents the control data while red represents epilepsy. The bold lines represent the mean while the shaded regions represent a 99% bootstrapped confidence interval. The bootstrapped null model produced via randomizing gives an average SSI of 0.23 ± 0.03 and is not shown here to increase visual clarity.

The same analysis performed in epilepsy reveals a region dependent alteration in SSI (Figure 3, red curves). There is a large decrease in SSI for all features in the mEC, indicating a loss of the constraint exerted by global oscillatory states on the selection of possible substates, again regardless to the chosen *k’*s. In contrast, there is no such large loss of brain state specificity in CA1, in particular no change for sharing. We conclude that the substate distribution becomes “disordered”, i.e., a large proportion of substates now occur during both THE and SO in the mEC in epilepsy. In contrast, CA1 retains the brain state specificity of the distribution of substates. The alteration of brain state-specificity of firing, sharing and storage substates is therefore brain region dependent in epilepsy.

### Computing hubs are more numerous but less substate-specific in the mEC in epilepsy

Within each substate/feature we extract computing hub neurons, i.e., neurons with on average, exceptionally high firing, storage or sharing values with regard to the substate (see Methods). As previously discussed in Clawson et al. (2019), it is important to stress that different substates are associated to different sets of hubs and that a neuron acting as firing, storage or sharing hub in a given substate will not necessarily do so in another substate. So, while the fraction of neurons being hub in a given substate remains small, the fraction of neurons serving as hub at least in one substate is much larger, approaching ~40% on average. Figure 4A illustrates an example of the distribution of hubs (same recording as in Figure 2A).

**Figure 4.**
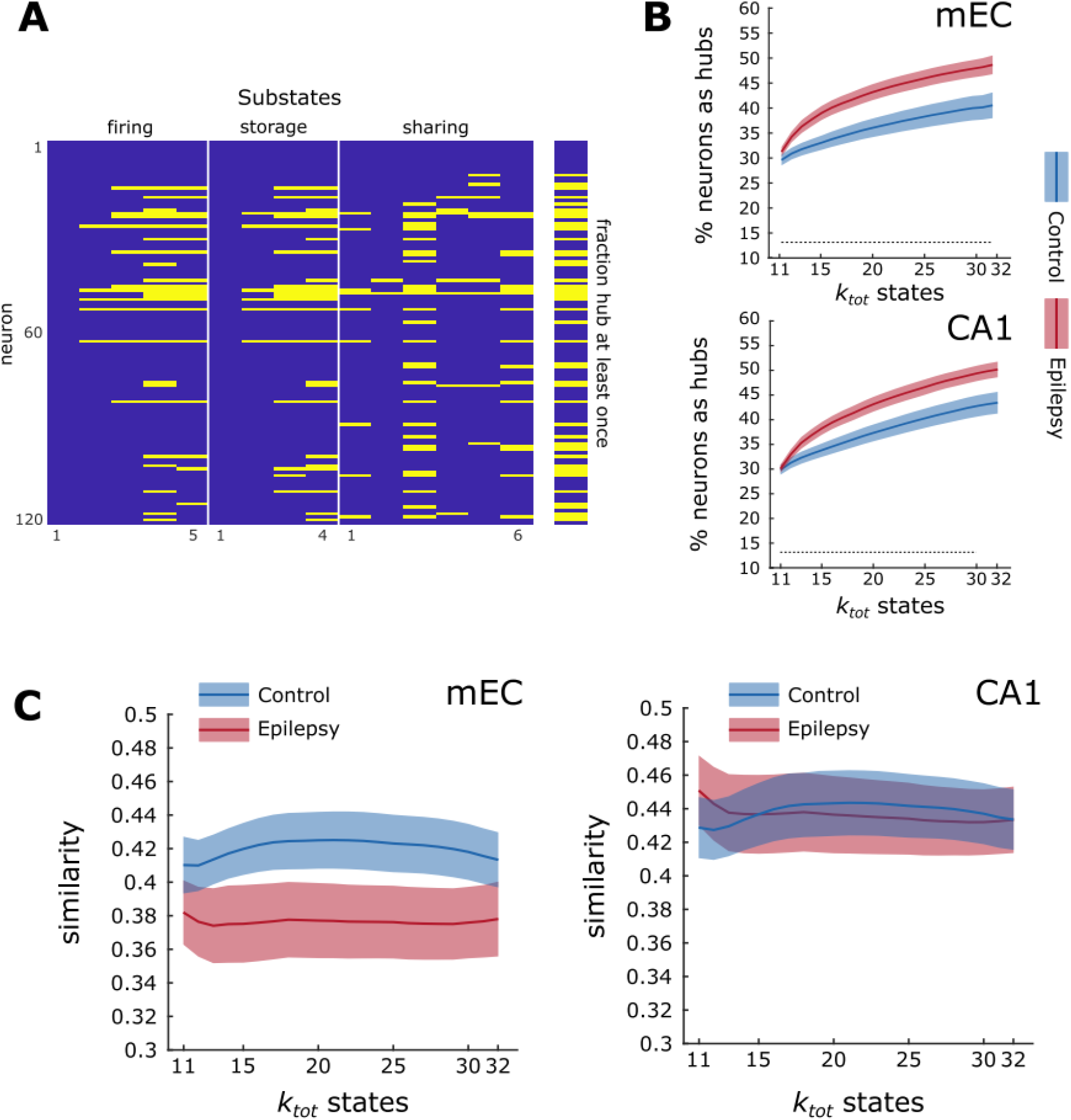
Computing Hubs and their distributions. (A) Example of computing hubs in the control mEC extracted from a given state table. The y axis is unsorted neuron label, and the x axis shows the substates for firing (5), storage (4) and sharing (6) features. A yellow bar indicates that the given neuron is a computational hub during a substate. On the right is a summed version of the graph on the left, visually showing the fraction of neurons that are a hub at least once (40%). (B) The percentage of neurons that are hubs at least once is increased in epilepsy independently of *k_tot_*. The grey dotted line represents the mean of the shuffled, null model. (C) The similarity index plotted as a function of *k_tot_*. The hubs become less substate-specific in the mEC in epilepsy. Blue and red are for control and epilepsy data, respectively. The bold lines are the mean, and the shaded regions are the 99% bootstrapped confidence interval.

In control animals, the percentage of hubs increases with *k_tot_* in both mEC and CA1 (Figure 4B), which is expected due to the arbitrary over-clustering as *k* increases. We observe furthermore that the percentage of neurons serving as hubs at least once is significantly increased in epilepsy, by 5% in the mEC and 2.5% in CA1 (Figure 4B). This result is in agreement with the increase in substate contrast found in epilepsy: more neurons are more contrasted and therefore are detected as hubs. Note that, for both control and epilepsy, the percentage of neurons marked as hubs is significantly larger as compared to randomized state tables (grey dotted lines in Figure 4B), confirming that the emergence of hubs is a direct fingerprint of the existence of well distinct substates.

Figure 4A also shows that some computing hubs are shared by different substates, while others are specific to one substate/one feature. In order to assess how substate-specific the computing hubs are, we use a measure of *similarity* (see Methods). A null value indicates that every substate has a unique hub set with no overlap between substates while a 1 value means that all substates have an identical distribution. Figure 4C shows that, in control animals, a majority of hubs tend to be substate-specific (similarity < 0.5). In CA1, the distribution of hubs is less substate-specific than in the mEC (higher similarity). In epilepsy, the distribution of hubs does not change in CA1, while hubs become significantly more substate-specific in the mEC. In other words, the status of being hub is for a mEC neuron less stable in epilepsy than in control animals.

We conclude that, in epilepsy, the mEC and CA1 display an increase in the number of neurons labeled as hubs at least once, and that the substate-specificity of hubs is increased in the mEC. Taken together, these two findings suggest a more hectic and random-like emergence of computing hubs in epilepsy as compared to control, albeit expressed in different ways; in mEC there are more hubs that are simultaneously more specific than control and in CA1 there are more hubs while staying the same, indicating a possible ‘shuffling’ of hubs. We believe this also further confirms that alterations in information processing are brain-region dependent.

### Alterations in the core-periphery organization of CA1 computing hubs in epilepsy

The partners from whom a given neuron receives or to whom it sends information are continuously changing (Clawson et al., 2019). At each time step, the instantaneous sharing networks can be seen as having a dynamic core-periphery structure (Pedreschi et al., 2020), with a core of tightly integrated neurons, surrounded by lightly connected periphery neurons. Two key measures of the core-periphery structure are the coreness, how central or well-integrated within a dense subnetwork – how “core”– a given neuron is, and the Jaccard index, a measure indicating how similar (or, conversely, liquid) the connections are between the recorded neurons between two time steps. We find that average coreness and the overall coreness distribution shapes are not significantly changed in epilepsy for either mEC or CA1 (Fig S5). Thus, the core-periphery architecture of information sharing networks within every substate is preserved in epilepsy. However, during the SO state, the average Jaccard values in CA1 are significantly decreased in epilepsy as compared to control (Fig S5). Thus, in CA1 there is enhanced connectivity variance and more volatile recruitment of neurons in the core.

### Assessing substate sequences

The analysis of individual features (firing, storage and sharing) revealed brain state- and brain region-dependent alterations in epilepsy. We now focus on a more integrated view of the informational patterns, in which we consider both the simultaneity of the ongoing types of patterns and their articulation in sequences along time. We perform this higher-level exploration using the notion of information processing states (IPS), driven by the idea of symbolization, as shown in Figure 2A (Porta et al., 2015). From each analysis time window, we generate a four-letter word, with the letters representing the substate labels of the global state, firing, storage and sharing features measured in this time window (see Methods). When the analysis window is shifted by 1 s, another word is obtained, which is identical to the previous one if the substate does not change. This procedure allows us to reduce the description of the complex simultaneous variations of firing, storage and sharing patterns within the neuronal population to simple strings of symbolic *words*, a sort of “neuronal language” built of sequences of possible words in a dictionary. We can then assess how the properties of these strings are modified in epilepsy at the level of their dictionary and syntax.

We defined all possible state tables generated through our k-means procedure as a *library* (Fig 2). Now, as tables are considered as a sequence of words, we define the sequence of words generated as a *book*. The number of letters, and therefore the number of words, depend upon *k_tot_*. As a result, we label our differently generated books by *k_tot_*. All 512 possible books per recording are grouped together to form a *library*. For each library, we build two sister libraries for comparison: one in which we sort every book internally to be highly ordered, and one in which we randomize every book internally to be highly disordered (see Methods). Using this word/book analogy, we begin to explore the organization of the language of the information processing contained in the books held within the library – What words are expressed? Is there a syntax, or organizational rules? And how does epilepsy change these measures?

### Impoverishment of the Dictionary in the mEC in Epilepsy

For each *k_tot_*, there is a fixed number of potential words that can be generated and possibly appear within the associated book (see cartoon in Figure 5A). As in any language, only a fraction of all possible words is expressed. For each book, we measure the used dictionary fraction, or *relative dictionary* (see Methods). Figure 5A illustrates two end cases. The low relative dictionary (left) uses a small number of expressed words, while the high relative dictionary (right) uses a much richer vocabulary, wherein almost all of the potential dictionary is expressed. While the measure of relative dictionary in and of itself is informative, it is difficult to use such a measure to assess meaningful changes (i.e., before control and epilepsy) without having comparative baselines. Therefore, we compute not only the relative dictionary of our libraries, but also that of the ordered and random sister libraries (which correspond to the null hypotheses of order and disorder in the ‘language’ of the book, respectively). We then apply a linear transformation to the relative dictionary measure, resulting in 0 representing the relative dictionary measure of ordered books, and a value of 1 representing a relative dictionary measure identical to that of randomized books. Such a normalized relative dictionary measure tracks not only the richness of the used dictionary but also its position between order and disorder.

**Figure 5.**
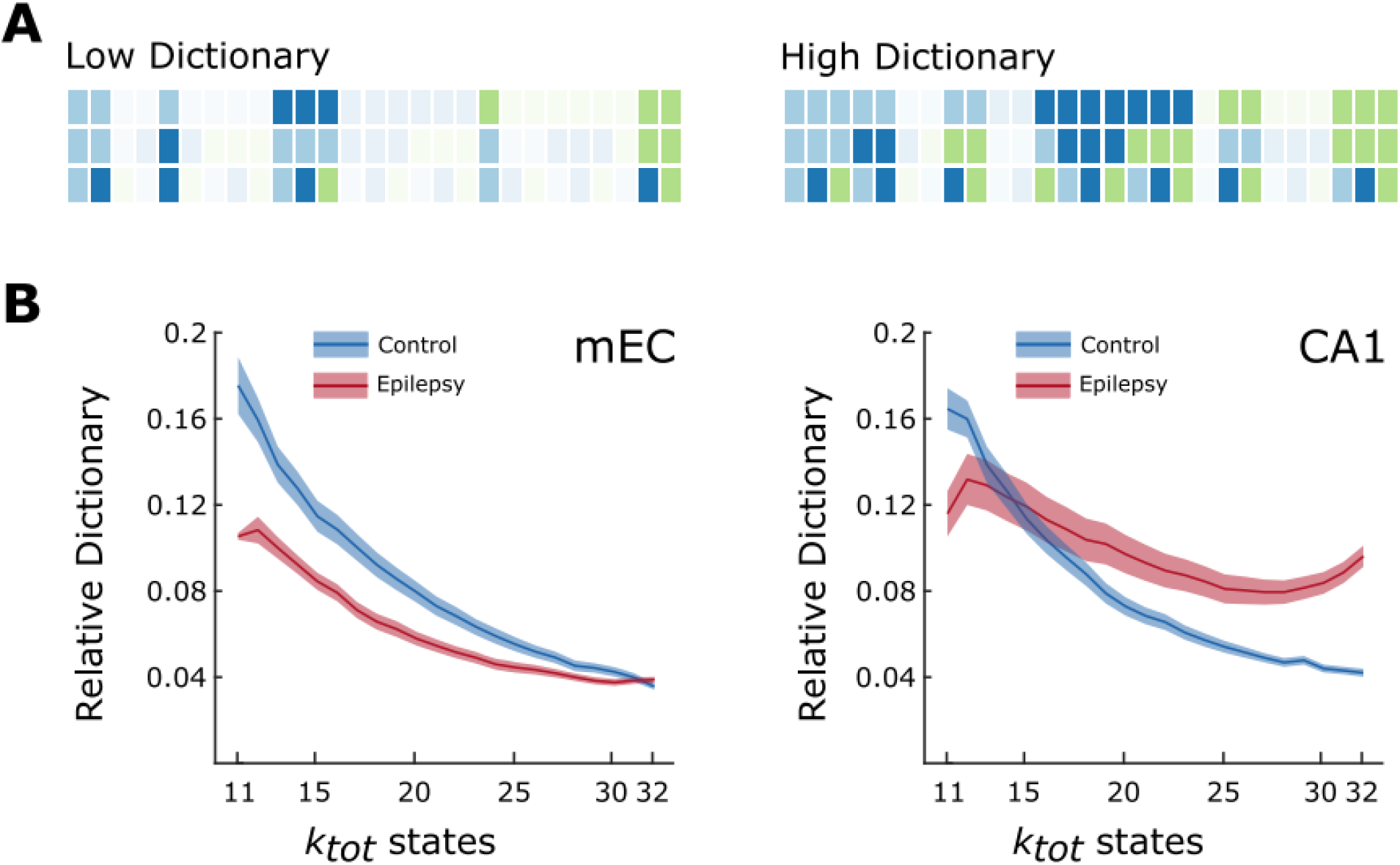
Relative dictionaries within the libraries. (A) Fictional cartoons representing two extremes for the measure of relative dictionary. Each row represents a feature (firing, storage, sharing); for simplicity we do not take into account the brain states (THE and SO). We consider three substates (light blue, dark blue, green) per feature (using the same color code for simplicity), which makes a total of 3^3^ = 27 words (the representation is similar to counting in base 3 with color, increasing from left to right). Words that are not observed are shaded. A low relative dictionary (left) contains a low fraction of all possible words, while a high relative dictionary (right) contains a high fraction. (B) Relative dictionary values as a function of *k_tot_*. As expected, the fraction of words used in control decreases as the number of possible words increases. The relative dictionaries are similar in mEC and CA1 in controls. There is a marked decrease in the relative dictionary in the mEC in epilepsy. In CA1, the relative dictionary in increased or decreased as compared to control as a function of *k_tot_*. Blue is representative of control data and red is representative of epileptic data. The bold lines are the mean, and the shaded regions are the 99% bootstrapped confidence interval.

Figure 5B shows that for both the mEC and CA1 in control and epilepsy conditions, the normalized relative dictionaries lie much closer to 0 than to 1, meaning that their relative dictionaries are much more similar to a system with organization that is ordered than disordered. In epilepsy, the relative dictionary is reduced with respect to control in the mEC (Figure 5B). Thus, the dictionary of state dynamics language seems impoverished in the mEC in epilepsy. There is also a reduction in CA1, but only for books with low *k_tot_* values whereas it is increased for *k_tot_* > 15. This ‘crossing’ of control and epileptic near *k_tot_* = 15 may be potentially explained by the strength of clustering for sharing features (Fig S3). Contrary to all features, there exists only a small window of *k* for sharing in CA1 in which k_means clusters the feature better than a null model. Therefore, dictionaries made with poor clustering may drive the dictionary too high for low values of *k*. This is the first instance for which the generic rule that results should be independent of the choice of k, fails. However, this characterization of dictionaries further demonstrates that the alterations are brain region dependent.

The relative dictionary provides important information about the words, but not how words are organized in time. This is similar to the grammar, or syntax, of a traditional sentence. To analyze this syntax (how words are organized from one window not the next), we quantify the level of organization present in the state tables as a whole, i.e., the overall dynamics of a system moving though IPSs (Figure 2A).

### The syntax of substate sequences is less regular in epilepsy

Compressibility is a key property of an object as it represents the degree of internal order of the object. This is because any regularity within may be described by simply referencing its previous occurrence. Again, our state tables are bordered by two extreme cases: order and randomness (Figure 6A). An ordered table is dominated by a highly structured syntax, typically dominated by a lower dictionary and long periods of sustained words. Therefore, an ordered table is very compressible due to this internal order. A random table, on the other hand, typically contains an exceedingly high number of words, which follow each other in a disorderly (random) manner. This results in non-compressibility. A complex table is one that lies between those extremes. In order to characterize the complexity of the state tables, we compute a tailored form of description length complexity (Clawson et al., 2019; Rissanen, 1978), which is scaled to the sister libraries of order and disorder. Thus, in Figure 6B, 0 represents the complexity of the ordered library, something very compressible, while 1 represents the complexity of our disordered library, something very uncompressible (as shown in Figure 6A). In controls, the complexity is similar in mEC and CA1, close to an ordered table. In epilepsy, the complexity is significantly increased for all *k_tot_* values, while it is increased in the mEC at the high end of our library.

**Figure 6.**
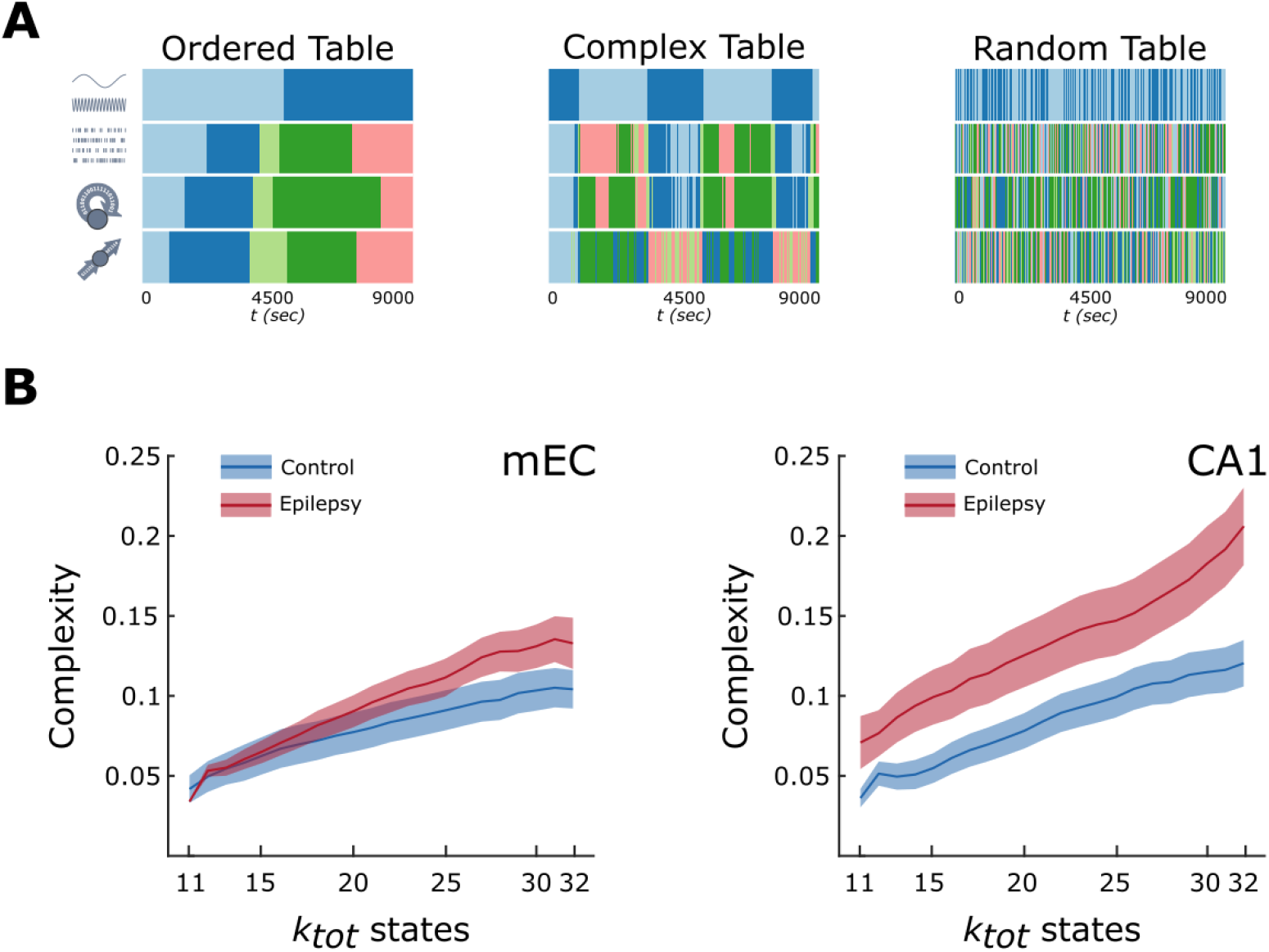
Order, disorder, and complexity. (A) Examples of state tables, similar to that of Fig 2A, from the mEC showing the two extremes of order and disorder as well as one of the possible state tables taken from the state table library. (B) Complexity values for both the mEC and CA1 as a function of *k_tot_*. The complexity is similar in mEC and CA1 in controls. In epilepsy, the complexity is largely increased in CA1, and only for large *k_tot_* values for mEC. Blue is representative of control data and red is representative of epileptic data. The bold lines are the mean, and the shaded regions are the 99% bootstrapped confidence interval.

Combining the results from Figures 5 & 6, we can propose the following interpretation. In CA1, the increase in complexity found in epilepsy, at least for books with sufficiently large *k_tot_*, can be explained, at least in part, by an enriched dictionary, since enrichment of the relative dictionary positively correlate with complexity (Clawson et al., 2019). In the mEC, the relative dictionary decreases while the complexity mildly increases. Thus, mEC books have a less regular syntax despite being constructed out of a lesser number of words.

## Discussion

This study provides evidence that epileptic conditions alter information processing in its simplest sense, the primitive storage and sharing operations as we introduce here, in a brain-region dependent manner. As these basic processes are necessarily involved in a variety of neural computations, their alterations may indirectly impact numerous cognitive functions.

The main limitation to our study is that it is made under anesthesia, versus for example, goal-directed behavior to assess cognitive function. The type of analysis we performed is powerful as it allows unraveling basic properties of information processing without needing to know which computations are ongoing. However, it requires long-duration, stable recordings with large state sampling to obtain enough data points to perform reliable statistics. We did not record during natural sleep, because seizures and interictal spikes (which would act as strong confounding factors) mostly occur during the light phase, while they do not occur under anesthesia. However, a similar type of analysis performed in control animals led to similar results during sleep and anesthesia (Clawson et al., 2019), suggesting that the anesthesia procedure we use does not significantly alter core information dynamics.

We refer to the elementary information storage and sharing operations as primitive (or low level) information processing operations, as we consider them as fundamental building blocks within an algorithm to reach an end condition (like a function), similar to the “algorithmic level”, introduced by Marr & Poggio (1977). Algorithm is used here in its most generic meaning, as we do not claim that the brain is analogous to a computer. Such primitive processing operations, as we define them, represent nothing else than the emergent “informational effect” of very concrete neurophysiological phenomena. Storage and sharing of information directly derive from auto- and cross-correlations in firing, which widely vary in neuronal populations (Schneidman et al., 2006), and can be directly measured from spiking activity of neurons. Other primitive processing operations exist, such as information transfer (Palmigiano et al., 2017; Schreiber, 2000) or information modification (Lizier et al., 2013; Wibral et al., 2017). Our recordings and choice of a time-resolved approach do not provide enough data to track these more sophisticated operations. However, the processing functions of storage and sharing are especially important as they represent statistical measures of information flow in time, and spacetime, respectively.

We show that primitive information processes are organized in temporal sequences of information processing substates (IPSs), which are extracted via a cluster analysis. We have used a non-biased approach, spanning many possible combinations of numbers of clusters. The fact that most results are independent from the choice of the number of clusters provides a strong argument for the genericity of our conclusions. With this approach, we demonstrate a degradation of complexity due to enhanced randomness in epilepsy. This conclusion stems from the convergence of complementary analyses. First, storage and sharing hubs are less robust, waxing and waning in a more erratic manner across substates and the recruitment of neurons into the integrated core of sharing networks is more volatile. Second, average storage and sharing strength are more similar between brain states, and this “dedifferentiation” occurs despite the higher contrast between substates. Third, the state specificity of IPS is reduced, i.e., many IPSs are now redundant between THE or SO. Together, these results imply that a change in brain state is no longer associated to strong specificity in information processing. Fourth, freed from the constraint of being strongly state-specific, the relative dictionary in epilepsy could, in principle, be increased. However, mEC has a decreased relative dictionary, which instead implies an ability to form unique IPSs. Yet, the description complexity of IPS sequences tends to be larger in epilepsy than control. In other words, IPS sequences have a less regular syntax despite being assembled out of less unique words.

The IPS dynamics of CA1 show, in general, less alterations than that of mEC. The fact that information processing is affected in brain region-dependent manner is an important result. The mEC and CA1 have distinct cytoarchitectures and different fates following an epileptogenic insult. Most striking is the loss of layer 3 in the mEC, and the injury of many pyramidal cells and interneurons in the CA1 region (Curia et al., 2008). It is not possible to assign a given alteration in information processing to a given morpho-functional changes in the mEC or CA1. Global brains states (THE and SO) and IPSs are emergent properties. Any change in any brain region can potentially affect neuronal dynamics anywhere from the local to the global scale. Therefore, the morpho-functional alterations in mEC or CA1 may contribute to any combination of local and global changes. However, changes in terms of information processing do not necessarily have to be homogenous across brain regions. In fact, brain region-specific modifications are expected as each region is embedded in different functional networks. How these brain-region specific changes contribute to comorbidities (such as cognitive deficit, anxiety, and depression) remain to be determined.

Our measure of complexity is that of compressibility, accounting for the internal structure, i.e., how internally ordered are IPS syntaxes. Any change in this internal organization would thus imply an underlying change in algorithmic operation, resulting in different computation in control and epilepsy conditions. Our measure of complexity does not allow distinguishing beween an increase in processing versus an increase in noise, as complexity would grow in both cases. Other measures can be used, but they would require more data (Crutchfield, 2011). However, in CA1, books with large *k_tot_* have an increased, rather than decreased dictionary size, which may explain the strong increase in sequence complexity. It is not clear, however, that this dictionary increase is a positive factor as it may reflect a more irregular IPS selection, with rare IPSs indicating dysfunction in IPS sequential production. Another possibility is that boosted IPS sequence complexity in CA1 and, at a lesser extent, mEC is a compensatory mechanism to generate a more sophisticated syntax to compensate for other shortages, such as reduced hub stability and degraded state-specificity of IPS.

In a biological context, the algorithmic level change comes as a result of altered collective, spiking activity and could lead to an entirely different expression of higher-level behavior, such as cognition. However, the question of whether this increase of complexity (decrease of internal order) observed in epilepsy is the source of cognitive deficits or not remains ultimately open. It has been theorized that “biological systems manipulate spatial and temporal structure to produce order – low variance – at local scales” in an effort to adapt and survive (Flack, 2019). Therefore, if networks are still functional in epilepsy conditions, are these manipulations now less effective? Or is the resulting low variance order now too difficult to sustain due to a combination of physiological and functional changes? These issues remain to be addressed. Nevertheless, the approaches presented here introduce valuable insight into aspects of the collective behavior of neural populations, and provide a quantitative framework to answer such questions.

In conclusion, the framework we introduce here to compare information processing between control and epilepsy, can be generalized to neurological disorders. Since most, if not all, of the latter, including migraine, Alzheimer’s disease, and Parkinson’s disease are associated with co-morbidities, it will be particularly interesting to determine whether information processing at the algorithmic level is also affected in these disorders. Following the principle of degeneracy (Prinz et al., 2004), very different structural alterations, which characterize different neurological disorders, may produce similar alterations in information processing, providing an explanation for the commonalities of co-morbidities across different disorders.

## Methods

### Ethics

All experiments were conducted in accordance with Aix-Marseille Université and Inserm Institutional Animal Care and Use Committee guidelines. The protocol was approved by the French Ministry of National Education, Superior Teaching, and Research, under the authorization number 01451-02. All surgical procedures were performed under anesthesia and every effort was made to minimize suffering and maximize the animals’ wellbeing from their arrival to their death. All the animals were housed in pairs in large cages with minimal enrichment, food and water at libitum, in a room with controlled environment (temperature: 22 ± 1 °C; 12 h light/dark schedule with lights off at 8:00 pm; hygrometry: 55%; ventilation: 15-20 vol/h).

### Data information

We use in this work a portion of the data (5 out of 7 original experiments) initially published by Clawson et al. 2019 as control data, which includes local field potentials (LFPs) and single-unit recordings obtained from the dorsomedial entorhinal cortex (mEC) and the dorsal hippocampus (HPC) of anesthetized rats. Six recordings are original data, which includes LFPs and single-units recorded in the mEC and HPC recorded simultaneously under anesthesia in epileptic condition. See Figures S1 for details on recordings, number of cells, and layers recorded.

### Epilepsy model and surgery

We induced status epilepticus (SE) on 6 male Wistar (250–400 g; Charles Rivers) by a single intraperitoneal (IP) injection of pilocarpine (320 mg/kg; Sigma-Aldrich), one week after receiving the animals from the vendor. To reduce peripheral effects, rats were pre-treated with methyl-scopolamine (1 mg/kg, IP; Sigma-Aldrich) 30 min before the pilocarpine injection. SE was stopped by diazepam (10 mg/kg, IP, two doses within a 15-min interval) after 60 min. Then the animals were hydrated with saline (2 ml, IP, twice within 2 h) and fed with a porridge made of soaked pellets, until they resumed normal feeding behavior.

At least 8 weeks after SE induction, we performed acute recordings. Rats were first quickly placed in isoflurane (4% in 2l/min O_2_) and injected IP with urethane (1.5 g/kg) and ketamine/xylazine (20 and 2 mg/kg, IM), additional doses of ketamine/xylazine (2 and 0.2 mg/kg) being supplemented during the electrophysiological recordings. At all times the body temperature was monitored and kept constant with a heating pad. Heart rate, breathing rate, pulse distension, and arterial oxygen saturation were also monitored with an oximeter (MouseOX; StarrLife Sciences) during the duration of the experiment to ensure the stability of the anesthesia and monitor the vital constants. The head was fixed in a stereotaxic frame (Kopf) and the skull was exposed and cleaned. Two miniature stainless-steel screws driven into the skull above the cerebellum served as ground and reference electrodes. Two craniotomies were performed to reach the mEC and the CA1 field of the HPC, respectively: from bregma: −7.0 mm AP and +4.0 mm ML; and from bregma: −3.0 mm AP and +2.5 mm ML. We chose these coordinates to respect known anatomical and functional connectivity in the cortico-hippocampal circuitry (Witter et al., 1988; Witter et al., 1989). Two 32-site silicon probes (NeuroNexus) were mounted on a stereotaxic arm each. A H1×32-10mm-50-177 was lowered at 5.0-5.2 mm from the brain surface with a 20° angle to reach the dorso-medial portion of the mEC, and a H4×8-5mm-50-200-177 probe was lowered at 2.5 mm from the brain surface with a 20° angle to reach dorsal CA1. The on-line positioning of the probes was assisted by: the presence of unit activity in cell body layers and the reversal of theta ([3 6] Hz in anesthesia) oscillations when passing from layer 2 to 1 for the mEC probe, and the presence in *stratum pyramidale* either of unit activity and ripples (80-150 Hz) for the HPC probe. At the end of the recording, the animals were injected with a lethal dose of Pentobarbital Na (150mk/kg, i.p.) and perfused intracardially with 4% paraformaldehyde solution. We confirmed the position of the electrodes (DiIC18(3) (catalog #46804A, InterChim) was applied on the back of the probe before insertion) histologically on 40 μm Nissl-stained section as reported previously in detail (Ferraris et al., 2018; Quilichini et al., 2010). We used only experiments with appropriate position of the probe for analysis.

### Data collection and spike sorting

Extracellular signal recorded from the silicon probes was amplified (1000x), bandpass filtered (1 Hz to 5 kHz) and acquired continuously at 32 kHz with a 64-channel DigitalLynx (NeuraLynx) at 16-bit resolution. We preprocessed the raw data using a custom-developed suite of programs (Csicsvari et al., 1999). The signals were down-sampled to 1250 Hz for the local field potential (LFP) analysis. Spike sorting was performed automatically, using KLUSTAKWIK (http://klustakwik.sourceforge.net (Harris et al., 2000)), followed by manual adjustment of the clusters, with the help of auto-correlogram, cross-correlogram and spike waveform similarity matrix (KLUSTERS software package, http://klusters.source-forge.net (Hazan et al., 2006)). After spike sorting, we plotted the spike features of units as a function of time, and we discarded the units with signs of significant drift over the period of recording. Moreover, only units with clear refractory periods and well-defined cluster were included in the analyses (Harris et al., 2000). Recording sessions were divided into brain states of theta (THE) and slow oscillation (SO) periods using a visual selection from the ratios of the whitened power in the HPC LFP [3 6] Hz theta band and the power of the mEC LFP neighboring bands ([1 3] Hz and [7 14] Hz), and assisted by visual inspection of the raw traces (Ferraris et al., 2018; Quilichini et al., 2010). We then used band-averaged powers over the same frequency ranges of interest as features for the automated extraction of spectral states via unsupervised clustering, which confirmed our manual classification. We determined the layer assignment of the neurons from the approximate location of their soma relative to the recording sites (with the largest-amplitude unit corresponding to the putative location of the soma), the known distances between the recording sites, and the histological reconstruction of the recording electrode tracks. Animals were recorded for at least two hours in order to get few alternations of THE and SO episodes.

### Feature Computation

As in our previous work, for each region recorded we computed 4 main features from the electrophysiological data: global oscillatory band, neuronal firing sets, active information storage and the information sharing. We also keep the same sliding window paradigm where each feature is computed within a *10 second* window, and then the window is then moved forward in time *1 second,* which gives a *9 second* overlap. Therefore, when features are computed as described below, they are computed in this windowed fashion. The global oscillatory band features were computed by examining the LFP from both EC and CA1 and computing spectral power within 8 unequally sized frequency ranges (0–1.5 Hz, 1.5–2 Hz, 2–3 Hz, 3–5 Hz, 5–7 Hz, 7– 10 Hz, 10–23 Hz and 23–50 Hz), averaged over all channels within each of the recorded layers. Firing sets, active information storage, and the information sharing networks were all computed using a binarized raster built from the temporal labeling of spike firing (see Data Collection and Spike Sorting). Spiking data was binned using a *50 ms* bin; if a neuron fired within a given bin the output is a ‘1’, and if not, a ‘0’. This, for example would mean that a 2-hour recording would be transformed from a *7200 second × N* neuron matrix to a 7200000 *× N* neuron matrix that is composed solely of 0’s and 1’s. Firing sets were computed by computing the average firing density for each neuron within a window, and after these averages were compiled into time-dependent vectors. This resulting matrix is the *Firing Features*. Active information storage was computed by measuring the mutual information of a neuron’s binarized spike train between a given window and the window previous. What active information storage seeks to capture is the temporal ordering of individual spiking neurons, rather than capturing neurons that fire temporally close to one another (such as in the firing features). The resulting matrix is the *Storage Features*. Information sharing is computed by measuring the mutual information between a given neuron’s binarized spike train within a window and another neuron’s binarized spike train in the window previous. This process is iterated over all possible neuron pairs. Information sharing captures a similar metric to that of active information storage, although the key difference is that information sharing captures not just the temporal ordering, but the spatio-temporal ordering of spike timing, as it is computed across neuron pairs, rather than individual neurons. The resulting matrix is the *Information Sharing*. Although these measures have only been briefly described here, we suggest to the interested reader to examine the methods presented in our previous work [REF] where they have been rigorously defined.

### Feature-Based Substate Extraction

State extraction for each recording were also computed using the methods of our previous work, namely based around k-means clustering of each feature. The exception here, is we no longer choose a stable number of K clusters in k-means. Rather we cluster our 3 raster-based computed features (firing, storage, sharing) 3 separate times with K ranging from K = 3, 4,… 10. The function ‘kmeans’ was used from the default MATLAB toolbox. More information can be found on the Mathworks website. These K values were chosen as they represented a clustering range of too gross to too fine based on previous findings. *K* <= 2 would represent the same, or less, number of states as global states, which was previously established to be too small (Clawson et al., 2019). The clustering became too fine when K >= 10, wherein many substates only appeared for brief time periods, and never re-occurred. For each feature there are 8 different clustering results, done in an unsupervised manner 3 times to ensure that our results do not rely on single instance of clustering. This gave our analysis an opportunity to compute all metrics defined below over a robust range of K, ensuring that we can investigate how our substate stable metrics and results vary with arbitrarily too little or too many substates.

To compute the null model for substate extraction the process detailed above was repeated with the time stamps of all firing, storage and sharing jittered. This therefore retains the global mean and variance. Then, k-means was run on this jittered dataset 3 times, to produce 3 different clustering of the randomized dataset. These were not modified after this step and were used in any instances where a null model was needed (i.e. for silhouette and contrast).

### Substate Tables

Our main meta-object of study is a state table, a combination of our four features into a matrix (4 x number of windows). Table generation is an iterative process, as we have 8 possible substate configurations per feature. First, k = 3 in cluster attempt 1 for firing (FIRE _K3C1_), k = 3 in cluster attempt 1 for storage (STORE_K3C1_), and k = 3 in cluster attempt 1 for sharing (SHARE_K3C1_), are used in conjunction with the clustered spectral substates to form substate table 1 (Figure 2A).

Then, FIRE _K3C1_, STORE_K3C1_, and SHARE_K4C1_ are used in conjunction with the clustered spectral substates to form substate table 2. After, FIRE _K3C1_, STORE_K3C1_, and SHARE_K5C1_ used in conjunction with the clustered spectral substates to form substate table 3. This process continues such that all combinations of possible k values have been saved for a total of 512 different substate tables, with the final table having FIRE _K10C1_, STORE_K10C1_, and SHARE_K10C1_. It is important to note that all tables have the same spectral clustering, as the 2 substates of SO and THE are extremely robust as discussed above. This entire process is then repeated for each clustering attempt, resulting in 3 sets of our 512 substate tables for each region for each recording. Where applicable, all results are given as a function of total k states per table (i.e. for state table 1, there are 2 global states, 3 firing, 3 storage and 3 sharing for a total k_total_ = 11).

To produce the ordered tables for the ‘ordered’ null model, each substate table was sorted such that all substates with label ‘1’ appeared first, label ‘2’ was second, and so on and so forth. This can easily be achieved with the MATLAB function *sort*. Note that there is only one possible version of this type of ordering, and therefore the sample size for ordered tables is the same as recordings (n = 5 for control, n = 6 for epilepsy). To produce the randomized tables, substate labels were randomly permuted in time. For this process, we used bootstrapping to produce as 5000 randomizations to ensure the random null model was as strong as possible. To do this, 90% of each table was taken, randomly permuted and saved. These resulting tables were used as the random null model for relative dictionary and complexity seen in Figure 5 & 6.

### Contrast

To calculate contrast for a given feature we first calculate its global mean for each neuron (i.e., global mean firing per neuron). Here, ‘global’ refers to the entire recording. We then calculate the substate mean for each neuron by concatenating all periods of a given substate and calculating the mean across the ‘entire’ substate. The formula for contrast is then defined as the difference between the substate mean firing rate and the global mean firing rate, normalized by the global mean firing rate.

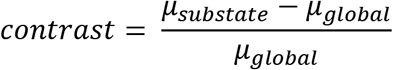

This allows the contrast to be either positive or negative. This process was done for all 3 features of firing, storage and sharing such that there are contrast values for each. This process was repeated for all possible clustering, therefore a contrast value per feature per *k*.

### Substate-Specificity

To compute the distribution of substates within periods of SO and THE we counted the number of times a substate appeared within a given epoch. Some substates exclusively appeared in only SO or THE, while others occurred in both. From these frequencies we estimated p(THE) and p(SO), i.e. the probability of a given substate occurring in either THE or SO, respectively. SSI is then:

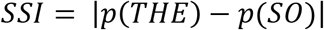

This equation results in SSI bound between 0 and 1, where 1 represents a state who exclusively occurs in either THE or SO and 0 represents a state that occurs equally in THE and SO.

### Hubs & Hub Stability

In this work we define a hub neuron in the same way as our previous work. Namely, for a given feature if a neuron’s activity within a given substate was higher than the 90^th^ percentile it was marked as a hub for the feature for that state. We compute hubs for every iteration of state table as defined above, such that we have a graph, or matrix, (see FIG 4A) for each state table. These matrices are Neuron × k_total_ where each entry is either a ‘0’ for non-hub or ‘1’ for hub. To compute how stable each of these matrices are as a function of k, we compute the normalized hamming distance of each matrix using the *pdist2* function in MATLAB but modified so that it gives a sense of how stable hubs are across states, where perfect similarity would result in a ‘1’, and no similarity at all would give a ‘0’.

### Coreness & Jaccard

The values for coreness & Jaccard were computed using the methods presented in Pedreschi et al. (2020). These were then analyzed using the same sliding window technique as presented in ‘Feature Computation’. After, periods of THE and SO were analyzed with similar techniques as that of Figure 1.

### Dictionary & Complexity

To compare sequences of substates of different types or in different regions we introduced a symbolic description of substate switching. With this description, each substate label acts as a letter symbol *s^(p)^*, where *(p)* can indicate firing, sharing or storage. For example, the firing features from the example substate table 1 [FIG 2A] would have the integer labels 1, 2, 3, and 4 (they can also arbitrarily be assigned letters as well, i.e. A, B, and C). We can therefore describe the temporal sequences of the visited substates of each feature as an ordered list of integers *s^(p)^(t).* Once substate labels are thought of as letters, we define the combination of firing, storage and sharing letters in each state table from a given window as 3 letter *words*. Using the formalism of linguistics, we can then compute the *dictionary,* or the number of words expressed, of a given recording within a region. We can also compute the *used dictionary fraction*, or the number of words found in the dictionary divided by the number of theoretically possible words given the number of substates per feature. For example, substate table 1 could have expressed 27 unique words. The used dictionary fraction was computed in an identical way to that of Clawson et al 2019. Specially, see ‘Complexity of substates sequences.

Using these methods, we compute the complexity of the sequences expressed using the notions of Kolmogorov-Chaitin complexity and minimum description length approaches (Crutchfield, 2011). While further discussion of method can be found here (Clawson et al., 2019) – the aspects of this complexity measure that is relevant for this work is that a random sequence of letters (and words) produces a higher complexity, while an ordered sequence of letters (and words) would produce a low complexity.

### Ordered & Random Substate Tables

To have relevant points of reference in our measures, each substate table was ordered and randomized. For the case of ordering, all substate labels for all features were sorted in ascending order which keeps the total lifetime of any state constant, while removing the temporal organization in an *ordered* fashion. In the case of randomization, all substate labels for all features were randomized 500 times, which again keeps the total lifetime of any state constant, while removing the temporal organization in a *random* fashion.

To compute the relative minimums and maximums for comparisons between order and random the MATLAB function ‘rescale’ was used. The minimums were computed using the average (of a given measure) of all ordered state tables for a given k_total_ and the maximums were computed using the average (of a given measure) of all random substate tables for a given k_total_.

### Plotting

Various tools were used for plotting. While mostly done via MATLAB, other tools were also used from ‘Moving Beyond p-values’ (Ho et al., 2019).

## Acknowledgements

P.P.Q. and A.G. performed and administered all surgery, implantation, and experimental recordings. P.P.Q, performed spike sorting, spectral analysis, and data pre-processing. W.C. and D.B. performed computational analysis. T.M. assisted with computational analysis and many of the supplementary materials. All authors designed the study and wrote the manuscript. W.C. is funded through the M-GATE program, who has received funding from the European Union’s Horizon 2020 research and innovation program under the Marie Skłodowska-Curie grant agreement No 765549. T.M was funded through Aix-Marseille Universite. P.P.Q. acknowledges support from FRM, FFRE and CURE Epilepsy Taking Flight Award. D.B. has benefitted for this work from support provided by the French Agence Nationale pour la Recherche (ERMUNDY, ANR-18-CE37-0014-02) and by the University of Strasbourg Institute for Advanced Study (USIAS) for a Fellowship, within the French national programme “Investment for the future” (IdEx-Unistra). C.B. is funded through ANR 19-CE14-0036-01. The funders had no role in study design, data collection and analysis, decision to publish, or preparation of the manuscript. The authors would like to thank Romain Goutagny and Anna Levina for scientific discussions and comments regarding this work.

## Data Availability

Partial data and codes can be found here: 10.5281/zenodo.4534369

Full codes, including figure generation as well as complete dataset are available upon request.

## Supplementary Figures

**S1.**
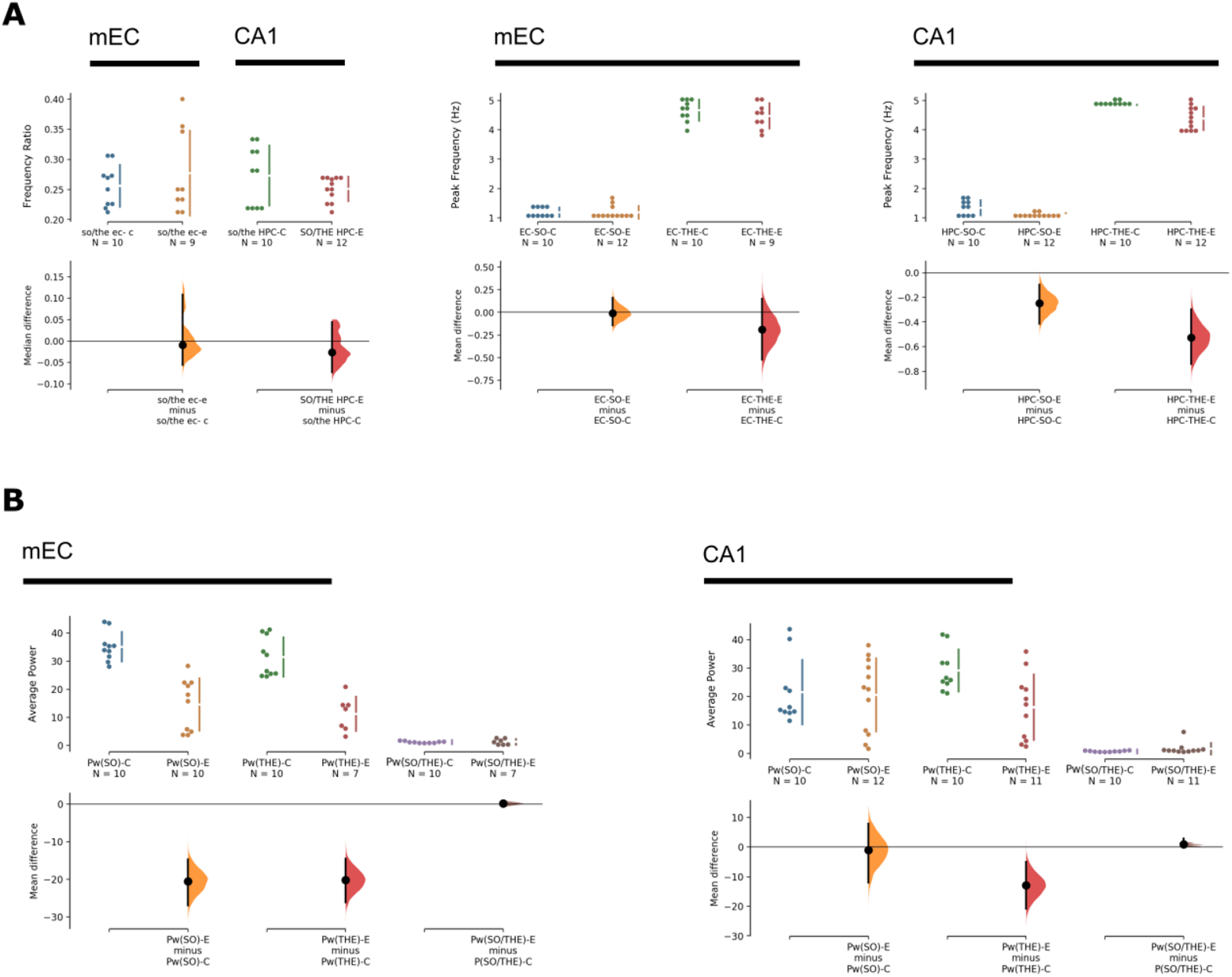
Frequency and Power Relationships in mEC and CA1 in control and epilepsy conditions. **(A)**(Far Left) A comparison of ratios between peak frequencies during periods of SO and THE in both control and epileptic conditions for mEC and CA1. (Middle and Right) Peak frequencies used in the previous graph for periods of SO and THE in control and epilepsy for mEC and CA1. There was a strong and smaller effect size on THE and SO peak frequency in CA1 in TLE, respectively. In these Cumming estimation plots, circles represent the mean, and all bars represent a 99% bootstrapped confidence interval. **(B)**The average power found in periods of SO and THE shown next to their ratio for both mEC and CA1. Note the strong effect size on THE and SO power in the mEC, and to a lesser extent on THE power in CA1. For all graphs, 5000 bootstrap samples were taken; the confidence interval is bias-corrected and accelerated.

**S2.**
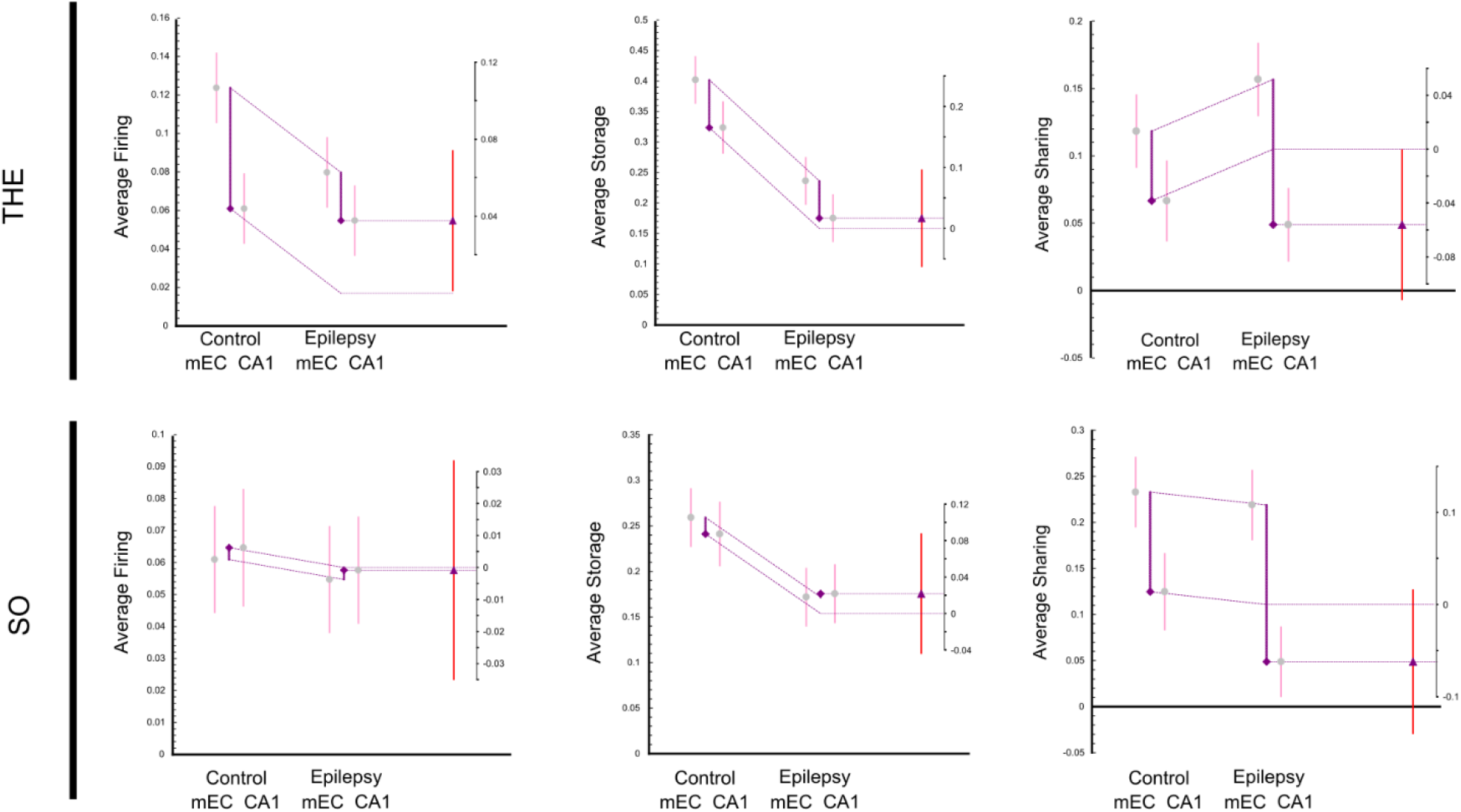
Average feature values presented as a function of mEC and CA1 in control and epilepsy conditions. The same graph as Figure 1D presented in an alternate format to highlight regional differences in control and epilepsy. The differences between mEC and CA1 during THE and SO are similar in control and epilepsy for average firing and average storage. The difference tends to increase for average sharing, but the effect size is consistent Circles and triangles represent the mean, and all bars represent a 99% bootstrapped confidence interval.

**S3.**
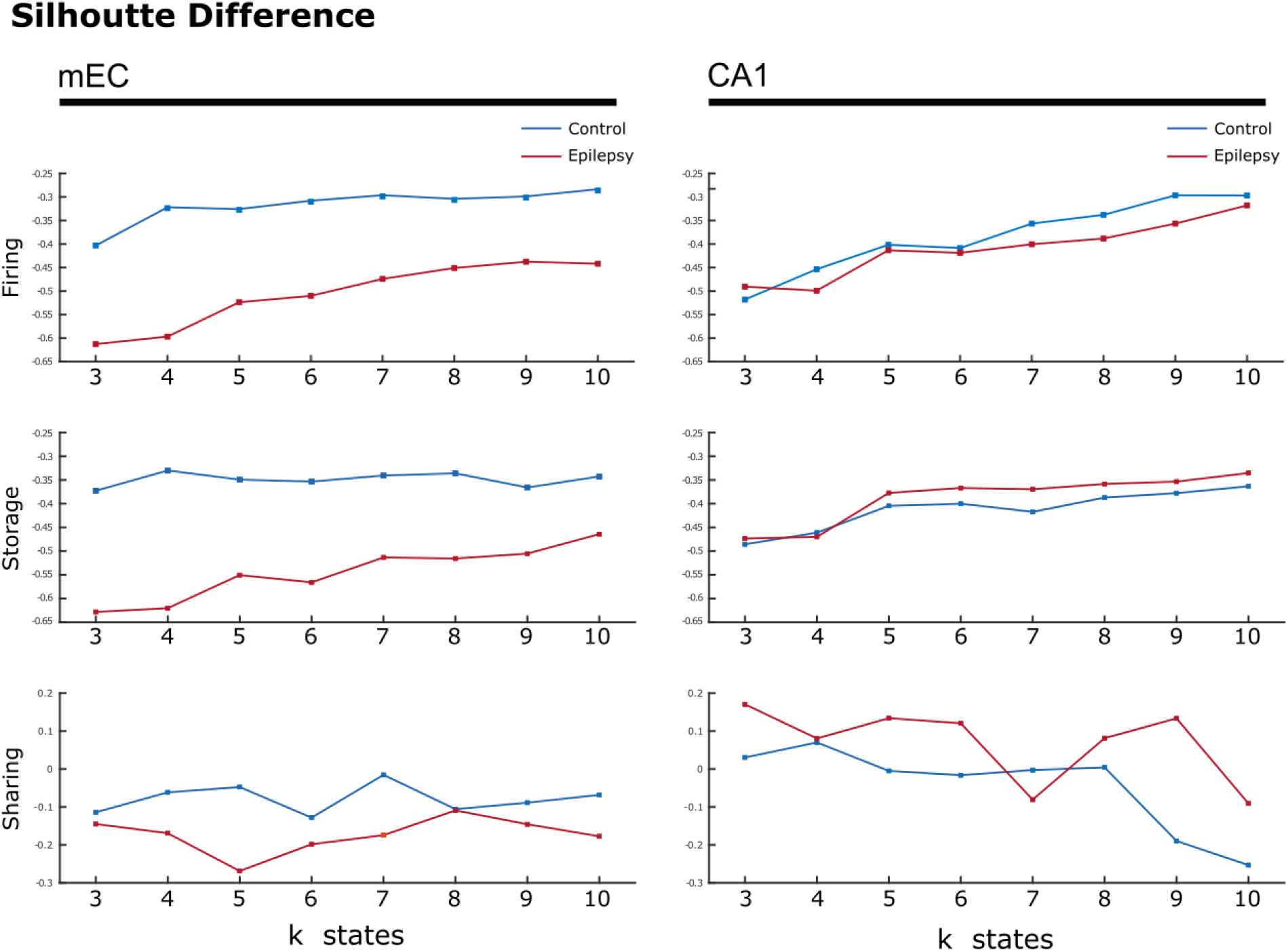
Null model with mean silhouette difference. The mean silhouette difference between a randomized null clustering model and the silhouettes found using k-means on non-shuffled data. Each point was calculated by computing mean silhouette values from a random selection of the randomized and normal clustering and taking the difference. This was done 500 times to produce error bars, but the error bars were so small that they appear to be squares on the graph. The blue line is representative of control data and the red line represents epilepsy data. There is a very large difference for firing and storage modalities from the null model for all *k* values in both CA1 and mEC in control and epilepsy conditions. Of special note is the sharing states found within CA1 (bottom right). We find that for both control and epilepsy conditions, our measure crosses 0 at k=5 and k=7, respectively, but fluctuates back above 0 until k=9 states in control and k=10 in epilepsy. This would indicate that the clustering only weakly holds in these intermediate values of k before not separating states better than a null, shuffled model up until the higher k values. Therefore, it may be that the states are either less definable in CA1 or, that on average there tend to be more states for sharing in both the control and epileptic states in CA1 and would therefore require higher k, on average.

**S4.**
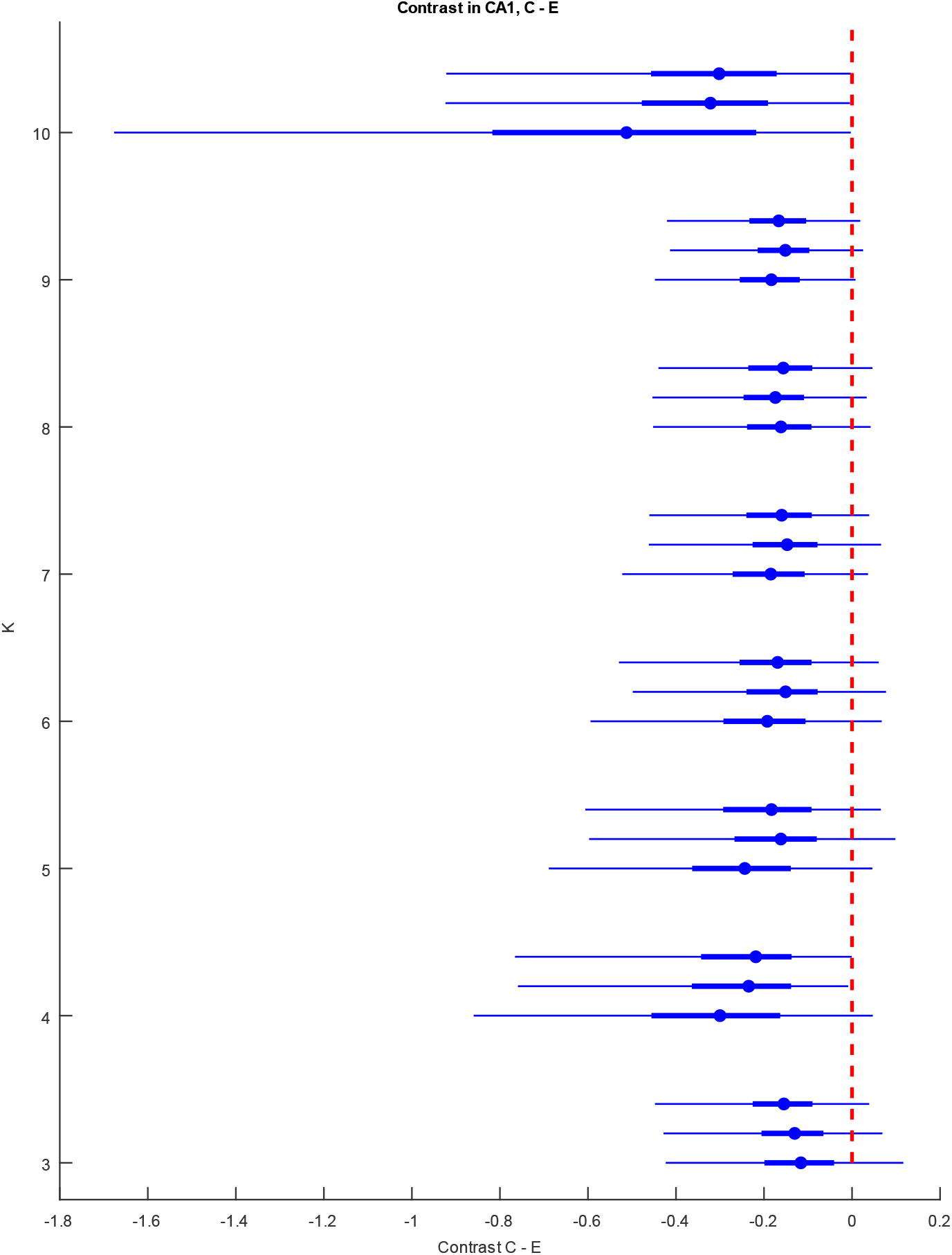
Contrast Values for Control vs Epilepsy in CA1. Average contrast difference between control and epilepsy is shown with respect to both feature and number of states, *k.* The circles represent the mean difference, the thick blue bars represent the 25-75% quantile and the thin blue bars represent the 1-99% quantile. The red dotted line is to add the null hypothesis line of no significant difference between control and epilepsy.

**S5.**
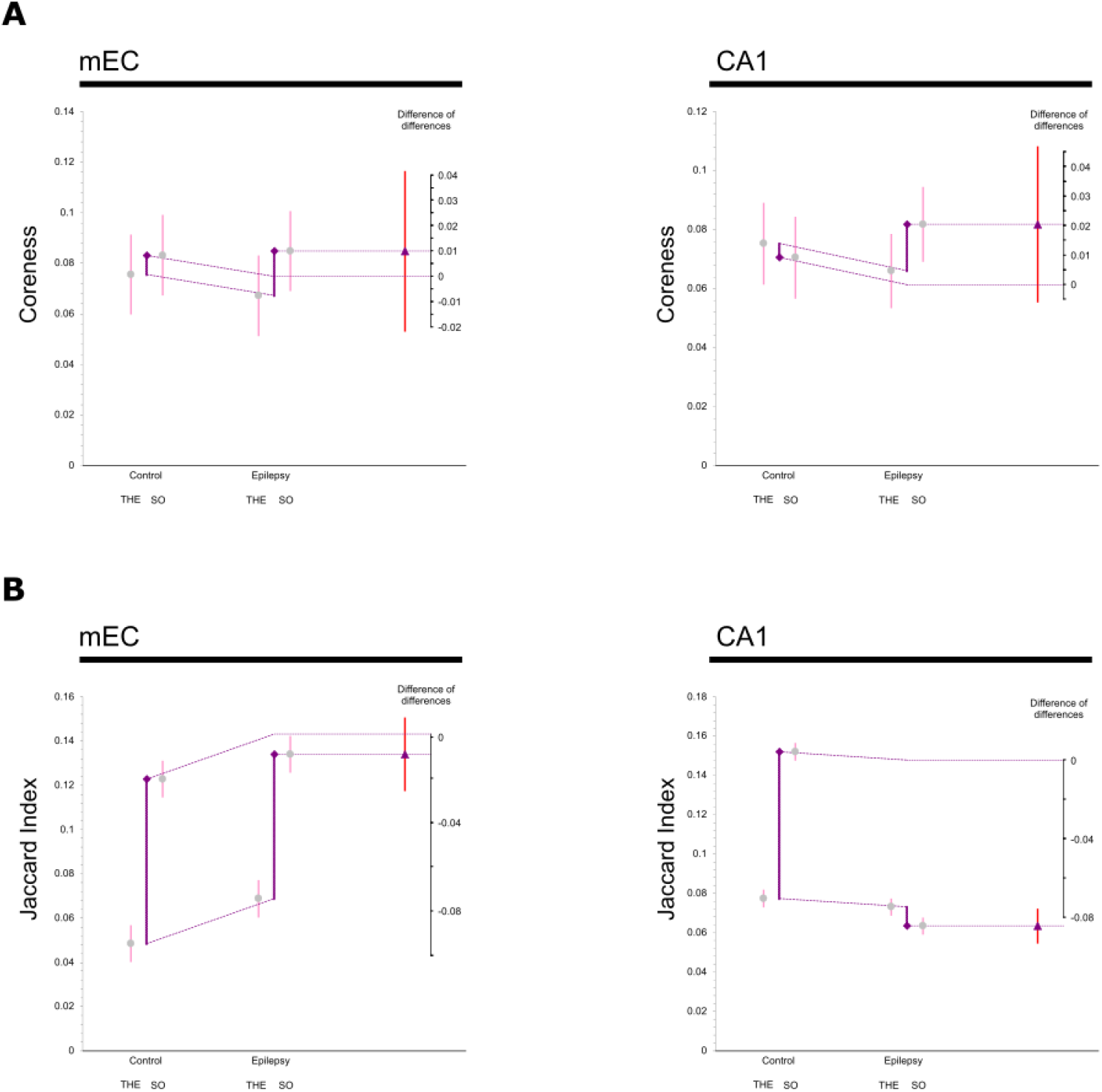
Coreness and Jaccard Values. Average values and difference of differences graphs for data features taken from sharing networks, found using the sharing feature, for both control and epileptic animals. Circles and triangles represent the mean, and all bars represent a 99% bootstrapped confidence interval. Note the very large effect size in the decrease of the Jaccard index in CA1 during SO. Accordingly, the brain state specificity of connectivity variance is lost. Significance is shown using the symbol (*) with their standard corresponding meaning (*, p<0.05; **, p<0.01; ***, p<0.001).

**ST1.** P value reporting: THE/SO unpaired mean difference. The p-value reported here is from a two-sided permutation t-test with CI intervals at 99%. 5000 bootstrap samples were taken; the confidence interval is bias-corrected and accelerated. The *P* value(s) reported are the likelihood(s) of observing the effect size(s) if the null hypothesis of zero difference is true. For each permutation *P* value, 5000 reshuffles of the control and test labels were performed. They are included here to satisfy a common requirement of scientific journals. (Ho et al., 2019)

**ST1 – P value reporting: THE/SO unpaired mean difference**
|  | mEC |  |  | CA1 |  |  |
| --- | --- | --- | --- | --- | --- | --- |
| **Firing** | mean difference | CI | P value | mean difference | CI | P value |
| Control | -0.0593 | [-0.0815, -0.0365] | 0 | -0.0475 | [-0.0823, -0.0162] | 0.0002 |
| Epilepsy | -0.0217 | [-0.0432, -2.55e-05] | 0.008 | 0.00082 | [-0.0226, 0.0263] | 0.00082 |
| **Storage** |  |  |  |  |  |  |
| Control | -0.133 | [-0.18, -0.0859] | 0 | -0.0828 | [-0.146, -0.0214] | 0.0002 |
| Epilepsy | -0.0584 | [-0.0933, -0.0224] | 0 | 0.0034 | [-0.0452, 0.0528] | 0.0034 |
| **Sharing** |  |  |  |  |  |  |
| Control | 0.109 | [0.0643, 0.16] | 0 | 0.0582 | [0.0369, 0.0814] | 0 |
| Epilepsy | 0.0535 | [-0.022, 0.128] | 0.0594 | 0.0285 | [0.0105, 0.0435] | 0 |

**ST2.**
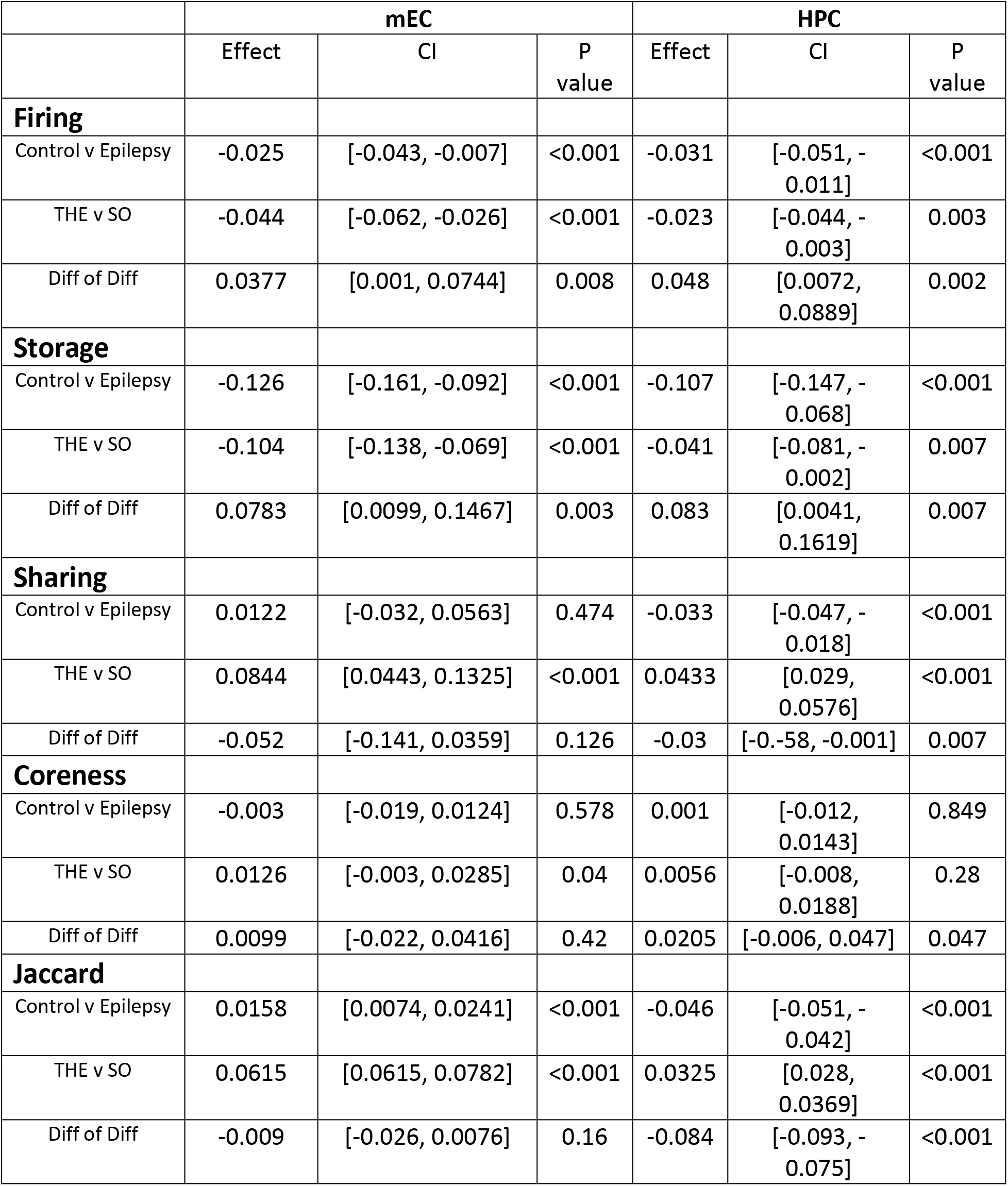
P value reporting: Difference of Difference Graphs.

## Bibliography

Barba, C., Barbati, G., Minotti, L., Hoffmann, D., & Kahane, P. (2007, Jul). Ictal clinical and scalp-EEG findings differentiating temporal lobe epilepsies from temporal ‘plus’ epilepsies. Brain, 130 (Pt 7), 1957–1967. https://doi.org/10.1093/brain/awm108

Bartolomei, F., Chauvel, P., & Wendling, F. (2008, Jul). Epileptogenicity of brain structures in human temporal lobe epilepsy: a quantified study from intracerebral EEG. Brain, 131 (Pt 7), 1818–1830. https://doi.org/10.1093/brain/awn111

Blumcke, I., Thom, M., Aronica, E., Armstrong, D. D., Bartolomei, F., Bernasconi, A., Bernasconi, N., Bien, C. G., Cendes, F., Coras, R., Cross, J. H., Jacques, T. S., Kahane, P., Mathern, G. W., Miyata, H., Moshe, S. L., Oz, B., Ozkara, C., Perucca, E., Sisodiya, S., Wiebe, S., & Spreafico, R. (2013, Jul). International consensus classification of hippocampal sclerosis in temporal lobe epilepsy: a Task Force report from the ILAE Commission on Diagnostic Methods. Epilepsia, 54 (7), 1315–1329. https://doi.org/10.1111/epi.12220

Calhoun, V. D., Miller, R., Pearlson, G., & Adali, T. (2014, Oct 22). The chronnectome: time-varying connectivity networks as the next frontier in fMRI data discovery. Neuron, 84 (2), 262–274. https://doi.org/10.1016/j.neuron.2014.10.015

Chauvière, L., Rafrafi, N., Thinus-Blanc, C., Bartolomei, F., Esclapez, M., & Bernard, C. (2009). Early deficits in spatial memory and theta rhythm in experimental temporal lobe epilepsy. Journal of Neuroscience, 29 (17), 5402–5410. https://doi.org/29/17/5402[pii]10.1523/JNEUROSCI.4699-08.2009

Clawson, W., Vicente, A. F., Ferraris, M., Bernard, C., Battaglia, D., & Quilichini, P. P. (2019, Jun). Computing hubs in the hippocampus and cortex. Sci Adv, 5 (6), eaax4843. https://doi.org/10.1126/sciadv.aax4843

Crutchfield, J. P. (2011). Between order and chaos. Nature Physics, 8 (1), 17–24. https://doi.org/10.1038/nphys2190

Csicsvari, J., Hirase, H., Czurko, A., Mamiya, A., & Buzsáki, G. (1999). Oscillatory coupling of hippocampal pyramidal cells and interneurons in the behaving Rat. Journal of Neuroscience, 19 (1), 274–287. https://doi.org/10.1523/JNEUROSCI.19-01-00274.1999

Curia, G., Longo, D., Biagini, G., Jones, R. S., & Avoli, M. (2008, Jul 30). The pilocarpine model of temporal lobe epilepsy. J Neurosci Methods, 172 (2), 143–157. https://doi.org/10.1016/j.jneumeth.2008.04.019

de Barros Lourenco, F. H., Marques, L. H. N., & de Araujo Filho, G. M. (2020, Jul). Electroencephalogram alterations associated with psychiatric disorders in temporal lobe epilepsy with mesial sclerosis: A systematic review. Epilepsy Behav, 108, 107100. https://doi.org/10.1016/j.yebeh.2020.107100

Ferraris, M., Ghestem, A., Vicente, A. F., Nallet-Khosrofian, L., Bernard, C., & Quilichini, P. P. (2018, Mar 21). The Nucleus Reuniens Controls Long-Range Hippocampo-Prefrontal Gamma Synchronization during Slow Oscillations. J Neurosci, 38 (12), 3026–3038. https://doi.org/10.1523/JNEUROSCI.3058-17.2018

Flack, J. C. (2019). Life’s Information Hierachy. Sante Fe Institute Press.

Harris, K. D., Henze, D. A., Csicsvari, J., Hirase, H., & Buzsáki, G. (2000). Accuracy of tetrode spike separation as determined by simultaneous intracellular and extracellular measurements. Journal of Neurophysiology, 84 (1), 401–414.

Hazan, L., Zugaro, M., & Buzsáki, G. (2006). Klusters, NeuroScope, NDManager: a free software suite for neurophysiological data processing and visualization. J Neurosci Methods, 155 (2), 207–216. https://doi.org/10.1016/j.jneumeth.2006.01.017

Hesdorffer, D. C. (2016, Jun). Comorbidity between neurological illness and psychiatric disorders. CNS Spectr, 21 (3), 230–238. https://doi.org/10.1017/S1092852915000929

Ho, J., Tumkaya, T., Aryal, S., Choi, H., & Claridge-Chang, A. (2019, Jul). Moving beyond P values: data analysis with estimation graphics. Nat Methods, 16 (7), 565–566. https://doi.org/10.1038/s41592-019-0470-3

Holmes, G. L. (2015, Jun). Cognitive impairment in epilepsy: the role of network abnormalities. Epileptic Disord, 17 (2), 101–116. https://doi.org/10.1684/epd.2015.0739

Inostroza, M., Brotons-Mas, J. R., Laurent, F., Cid, E., & de la Prida, L. M. (2013, Nov 6). Specific impairment of “what-where-when” episodic-like memory in experimental models of temporal lobe epilepsy. J Neurosci, 33 (45), 17749–17762. https://doi.org/10.1523/JNEUROSCI.0957-13.2013

Kirst, C., Timme, M., & Battaglia, D. (2016, Apr 12). Dynamic information routing in complex networks. Nat Commun, 7 (7), 11061. https://doi.org/10.1038/ncomms11061

Krishnan, V. (2020, Jul 14). Depression and Anxiety in the Epilepsies: from Bench to Bedside. Curr Neurol Neurosci Rep, 20 (9), 41. https://doi.org/10.1007/s11910-020-01065-z

Lenck-Santini, P. P., & Holmes, G. L. (2008, May 7). Altered phase precession and compression of temporal sequences by place cells in epileptic rats. J Neurosci, 28 (19), 5053–5062. https://doi.org/10.1523/JNEUROSCI.5024-07.2008

Lenck-Santini, P. P., & Scott, R. C. (2015, Sep 3). Mechanisms Responsible for Cognitive Impairment in Epilepsy. Cold Spring Harb Perspect Med, 5 (10). https://doi.org/10.1101/cshperspect.a022772

Lizier, J. T., Atay, F. M., & Jost, J. (2012, Aug). Information storage, loop motifs, and clustered structure in complex networks. Phys Rev E Stat Nonlin Soft Matter Phys, 86 (2 Pt 2), 026110. https://doi.org/10.1103/PhysRevE.86.026110

Lizier, J. T., Flecker, B., & Williams, P. L. (2013). Towards a Synergy-based Approach to Measuring Information Modification. 2013 Ieee Symposium on Artificial Life (Alife), 43–51. https://doi.org/10.1109/ALIFE.2013.6602430

Lopez-Pigozzi, D., Laurent, F., Brotons-Mas, J. R., Valderrama, M., Valero, M., Fernandez-Lamo, I., Cid, E., Gomez-Dominguez, D., Gal, B., & Menendez de la Prida, L. (2016, Nov-Dec). Altered Oscillatory Dynamics of CA1 Parvalbumin Basket Cells during Theta-Gamma Rhythmopathies of Temporal Lobe Epilepsy. eNeuro, 3 (6). https://doi.org/10.1523/ENEURO.0284-16.2016

Marr, D. C., & Poggio, T. (1977). From Understanding Computation to Understanding Neural Circuitry. Neurosciences Research Program Bulletin, 15 (3), 470–491. <Go to ISI>://WOS:A1977EH37300024

Palmigiano, A., Geisel, T., Wolf, F., & Battaglia, D. (2017, Jul). Flexible information routing by transient synchrony. Nat Neurosci, 20 (7), 1014–1022. https://doi.org/10.1038/nn.4569

Pedreschi, N., Bernard, C., Clawson, W., Quilichini, P., Barrat, A., & Battaglia, D. (2020). Dynamic core-periphery structure of information sharing networks in entorhinal cortex and hippocampus. Network Neuroscience, 1‒30. https://doi.org/10.1162/netn_a_00142

Porta, A., Baumert, M., Cysarz, D., & Wessel, N. (2015, Feb 13). Enhancing dynamical signatures of complex systems through symbolic computation. Philos Trans A Math Phys Eng Sci, 373 (2034). https://doi.org/10.1098/rsta.2014.0099

Prinz, A. A., Bucher, D., & Marder, E. (2004, Dec). Similar network activity from disparate circuit parameters. Nat Neurosci, 7 (12), 1345–1352. https://doi.org/10.1038/nn1352

Quilichini, P., Sirota, A., & Buzsáki, G. (2010). Intrinsic circuit organization and theta-gamma oscillation dynamics in the entorhinal cortex of the rat. J Neurosci, 30 (33), 11128–11142. https://doi.org/10.1523/JNEUROSCI.1327-10.2010

Quilichini, P. P., & Bernard, C. (2012). Brain state-dependent neuronal computation. Front Comput Neurosci, 6, 77. https://doi.org/10.3389/fncom.2012.00077

Rissanen, J. (1978). Modeling by shortest data description. Automatica, 14 (5), 465–471. https://doi.org/10.1016/0005-1098(78)90005-5

Rusina, E., Bernard, C., & Williamson, A. (2021). Kainic Acid Models of Temporal Lobe Epilepsy. eNeuro.

Schneidman, E., Berry, M. J., 2nd, Segev, R., & Bialek, W. (2006, Apr 20). Weak pairwise correlations imply strongly correlated network states in a neural population. Nature, 440 (7087), 1007–1012. https://doi.org/10.1038/nature04701

Schreiber, T. (2000, Jul 10). Measuring information transfer. Phys Rev Lett, 85 (2), 461–464. https://doi.org/10.1103/PhysRevLett.85.461

Scott, R. C., Menendez de la Prida, L., Mahoney, J. M., Kobow, K., Sankar, R., & de Curtis, M. (2018, Aug). WONOEP APPRAISAL: The many facets of epilepsy networks. Epilepsia, 59 (8), 1475–1483. https://doi.org/10.1111/epi.14503

Shannon, C. E. (1948). A Mathematical Theory of Communication. Bell System Technical Journal, 27 (3), 379–423. https://doi.org/10.1002/j.1538-7305.1948.tb01338.x

Suarez, L. M., Cid, E., Gal, B., Inostroza, M., Brotons-Mas, J. R., Gomez-Dominguez, D., de la Prida, L. M., & Solis, J. M. (2012). Systemic injection of kainic acid differently affects LTP magnitude depending on its epileptogenic efficiency. PLoS One, 7 (10), e48128. https://doi.org/10.1371/journal.pone.0048128

Tatum, W. O. t. (2012, Oct). Mesial temporal lobe epilepsy. J Clin Neurophysiol, 29 (5), 356–365. https://doi.org/10.1097/WNP.0b013e31826b3ab7

Valero, M., Averkin, R. G., Fernandez-Lamo, I., Aguilar, J., Lopez-Pigozzi, D., Brotons-Mas, J. R., Cid, E., Tamas, G., & Menendez de la Prida, L. (2017, Jun 21). Mechanisms for Selective Single-Cell Reactivation during Offline Sharp-Wave Ripples and Their Distortion by Fast Ripples. Neuron, 94 (6), 1234–1247 e1237. https://doi.org/10.1016/j.neuron.2017.05.032

Van de Ville, D., Britz, J., & Michel, C. M. (2010, Oct 19). EEG microstate sequences in healthy humans at rest reveal scale-free dynamics. Proc Natl Acad Sci U S A, 107 (42), 18179–18184. https://doi.org/10.1073/pnas.1007841107

Wibral, M., Lizier, J. T., Vogler, S., Priesemann, V., & Galuske, R. (2014). Local active information storage as a tool to understand distributed neural information processing. Front Neuroinform, 8, 1. https://doi.org/10.3389/fninf.2014.00001

Wibral, M., Priesemann, V., Kay, J. W., Lizier, J. T., & Phillips, W. A. (2017, Mar). Partial information decomposition as a unified approach to the specification of neural goal functions. Brain Cogn, 112, 25–38. https://doi.org/10.1016/j.bandc.2015.09.004

Witter, M. P., Griffioen, A. W., Jorritsma-Byham, B., & Krijnen, J. L. M. (1988). Entorhinal projections to the hippocampal CA1 region in the rat: An underestimated pathway. Neuroscience Letters, 85 (2), 193–198. https://doi.org/10.1016/0304-3940(88)90350-3

Witter, M. P., Groenewegen, H. J., Lopes da Silva, F. H., & Lohman, A. H. M. (1989). Functional organization of the extrinsic and intrinsic circuitry of the parahippocampal region. Progress in Neurobiology, 33 (3), 161–253. https://doi.org/10.1016/0301-0082(89)90009-9

